# Unveiling the small protein landscape of uropathogenic *E. coli*

**DOI:** 10.64898/2026.09.28.754932

**Authors:** Wenting Liu, Rick Gelhausen, Lisa Werland, Elisabetta Fiore, Florian C. Gruen, Eric Mersinger, Camila Cilveti, Timo Glatter, Cynthia M. Sharma, Rolf Backofen, Jing Yuan

## Abstract

Uropathogenic *Escherichia coli* (UPEC) is the predominant causative agent of urinary tract infections. The UPEC strain, CFT073, has a mosaic genome containing 60 genomic islands enriched in virulence genes. In this study, we explore the small proteome of CFT073 using ribosome profiling (Ribo-seq) coupled with RNA sequencing (RNA-seq), and reveal 338 translated, previously unannotated, putative small open reading frames (sORFs) with less than 100 codons. Among them, 78 sORFs are located in intergenic regions. Fifteen intergenic sORFs encode a predicted transmembrane helix (smemORFs), of which seven are hypothetical or have not been characterized in other *E. coli* strains. The majority of the remaining putative sORFs encodes uncharacterized or novel soluble proteins (ssolORFs). Considering the evolution of many sORFs is likely to be recent, functional sORFs are more likely to have conserved gene neighborhood in other genomes. Searching into the genomes of 10 other *E. coli* strains and four enterobacteria species, synteny analysis reveals that 10 of 15 putative smemORFs, and 27 of 63 putative ssolORFs, have a homologous sequence in a similar genomic context in at least one other genome, supporting their likelihood of encoding functional proteins. Further validation of putative sORFs using proteomics, cell-free protein synthesis, and mNeonGreen tagging provide strong evidence for the translation of nine predicted intergenic sORFs. Notably, four previously uncharacterized small membrane proteins have increased abundance under oxidative stress, suggesting a potential role in the stress response. Together, our findings reveal the small proteome of an extraintestinal pathogenic *E. coli* and provide a catalog of putative sORFs for further investigation.

## Introduction

Small open reading frames (sORFs) are historically unannotated and overlooked due to their small size, relatively low sequence conservation, and limitations of early gene prediction tools. Increasing evidence has shown that sORFs encode small proteins and play essential roles in diverse biological processes in all three domains of life (1–5). In prokaryotes, tens to hundreds of sORFs have been identified in different species (6–11). Functional characterizations reveal their interactions with larger proteins and the cell membrane, modulating their stability and function (12). For example, in *E. coli*, the small protein AcrZ (49aa) interacts with the bacterial RND transporter AcrB and modifies the substrate specificity of the drug efflux pump AcrAB-TolC (13, 14). The absence of AcrZ results in enhanced sensitivity of bacteria to specific antibiotics (13). In *Klebsiella pneumoniae*, disruption of the gene encoding the small protein MgrB (47 aa) is found to be the main mechanism responsible for the hyper-drug-resistant phenotype (15, 16). In humans, over 7000 sORFs have been identified, with several hundred small proteins validated. Some small proteins act as a major source of tumor-specific antigens, essential for immune surveillance, while others are involved in controlling development and morphogenesis (17, 18). Evolutionary analysis of sORFs and small proteins revealed that they are often acquired from *de novo* “gene birth” and are considered very young in the course of evolutionary history (19–22).

Next-generation sequencing of the complete set of RNA transcripts (RNA-seq) has provided accurate and comprehensive gene expression profiling in bacteria (23). It generates data confirming the transcription of sORFs with information on transcription start and stop sites. However, this approach does not provide evidence on the translation of sORFs nor the abundance of predicted small proteins in cells. The development of the ribosome profiling technique (Ribo-seq) complements the limitations of RNA-seq (24). Ribo-seq combines the next-generation sequencing technology with the isolation of actively translating ribosomes, allowing the determination of ribosome density on mRNAs by footprinting and providing direct evidence on the translation of sORFs. Compared to mass spectrometry-based proteomic approaches, Ribo-seq does not have requirement on specific protein sequences and has higher sensitivity for detecting small proteins. This approach has led to the discovery of small ORFs in diverse organisms, including many prokaryotes (6, 8, 9, 25–28).

The *E. coli* K-12 strain MG1655 was one of the first organisms for the systematic discovery of small proteins (29, 30). Approaches such as Ribo-seq and proteomics have revealed 150 new small proteins in this strain, expanding its small proteome beyond the number of proteins initially discovered (7, 31–34). Up to this point, the small proteome of MG1655 has 45 proteins (30% of the small proteome) containing a predicted transmembrane helix. More than 70% of the small proteome (108 out of 150) has been validated by Western blot, and some of these proteins have been further characterized functionally (7, 31–33). They are mostly small membrane proteins. In addition to AcrZ (49 aa) mentioned earlier, MgtS (31 aa) is another characterized small membrane protein. It is expressed under low magnesium conditions, binds to the magnesium transporter MgtA and cation-phosphate symporter PitA to improve cellular magnesium concentration (35, 36). PmrR (48 aa) is another membrane-bound small protein. Deletion of *pmrR* results in a growth defect in *E. coli* (37). In *Salmonella*, PmrR interacts with LpxT, an enzyme responsible for LPS modification, thus modifying the cell surface (38).

Uropathogenic *E. coli* (UPEC) strains are the primary cause of urinary tract infections, accounting for approximately 120 million infections developing worldwide each year (39). It was estimated that about 40% of women develop at least one UTI in their lifetime, and many of them experience recurrent UTIs annually (40, 41). Most UPEC cells originate in intestinal flora, enter the bladder via the urethra, cause infections in the bladder (cystitis) and, in severe cases, cause infections in the kidney (pyelonephritis) (42, 43). UPEC strains acquire virulence genes via horizontal gene transfer and mobile elements, enabling them to adapt to specific environments in the urinary tract and to infect hosts.

The UPEC strain CFT073 was isolated from a patient with severe pyelonephritis. It is studied extensively to understand pathogenicity, bacteria-host interaction, and antimicrobial resistance. The genome of CFT073 (about 5M bp in size) consists of 5490 annotated protein-coding genes in total and 60 genomic islands, which comprise about 15% of the genome (44). A range of virulence genes is located in these islands, including several genes encoding fimbrial adhesins, iron-uptake systems, hemolysin, and putative autotransporters (44). CFT073 shares 41.7% and 41.8% of total protein-coding genes with the *E. coli* lab strain MG1655 and the enterohaemorrhagic *E. coli* (EHEC) strain EDL933, respectively (44–46). The small proteome of *E. coli* MG1655 is among the best characterized, and that of EHEC is described as well (47). However, the small proteome of UPEC remained unknown. Here, we used ribosome profiling coupled with RNA sequencing and identified 78 sORFs in the intergenic regions of the CFT073 genome. Fifteen of the sORFs had homologues with annotated function in other genomes. The remaining sORFs encoded uncharacterized, hypothetical, and novel proteins. Genomic neighborhood analysis revealed 10 of 15 putative membrane protein-encoding sORFs, and 27 of 63 putative soluble protein-encoding sORFs, have homologous genomic context in at least one out of 13 bacterial genomes, supporting their likelihood of encoding functional proteins. Through proteomic analysis, cell-free protein synthesis, and mNeonGreen tagging approaches, we validated the translation of nine putative sORFs. Compared with other stress conditions (elevated temperature, high osmolarity, membrane perturbation, and nutrient limitation), oxidative stress led to an increased production of four previously uncharacterized small membrane proteins, suggesting a potential functional association with the oxidative stress response. The detailed information of identified sORFs (ssolORFs and smemORFs) are summarized in the supplementary file S1 and S2, respectively.

## Methods

### Strains and DNA templates

The uropathogenic *E. coli* strain CFT073 was used for this study. Linear DNA templates for the cell-free system were generated by overlapping PCR as described previously (48). A T7 promoter, a ribosome binding site and a transcription terminator sequence were included in the primers. The PCR products were then purified using a GeneJET PCR Purification Kit (Thermo Fisher Scientific). Primers used for producing the DNA templates for cell-free synthesis are listed in the supplementary Table S1.

### Construction of plasmids

For the expression of small proteins from their native promoters on a high copy plasmid containing the gene of mNeonGreen, first plasmid pLW73 was constructed from a pTrc99a backbone by removing the trc promoter and inserting a flexible linker and the gene sequence of mNeonGreen. For this, the gene of mNeonGreen was amplified and inserted into an amplified pTrc99a backbone without the trc promoter using Gibson assembly. Plasmids pLW74-pLW83 were then constructed based on pLW73. The native promoters and sORFs were amplified from the genome of E. coli CFT073 and inserted into an amplified pLW73 using Gibson assembly. Primers used for Gibson assembly and the resulting plasmids are listed in the supplementary Table S2 and S3, respectively.

### Sample preparation for ribosome profiling experiment

For the Ribo-seq experiment, *E. coli* UPEC CFT073 cells were grown aerobically in three biological replicates at 37°C in LB medium until they reached the early exponential phase (OD_600nm_ of 0.4). One hundred milliliters of the culture (equivalent to 40 OD_600_ units) were then harvested by fast-chilling as previously described (8, 9). Specifically, to block translation, 50 ml of the culture was transferred to cold 250 ml flasks and chilled for three minutes in an ice-water bath. The cells were transferred to 50 ml falcon tubes and centrifuged at 4,000 x g for 10 minutes at 4°C, before snap-freezing in liquid nitrogen. Prior to centrifugation, a sample of the culture was harvested for total RNA analysis and mixed with 0.2 volumes of STOP solution [5% buffer-saturated phenol (Roth) in 95% ethanol] and snap frozen in liquid nitrogen.

### Preparation of ribosome footprints

Ribosome profiling was performed generally as described previously with some modifications (6, 8–10, 49, 50). To lyse the cells, the bacterial cell pellet was resuspended in 1 ml of cold lysis buffer [100 mM NH_4_Cl, 10 mM MgCl_2_, 20 mM Tris-HCl pH 7.5, 0.4% Triton X-100, 1 mM PMSF, 150 U of DNase I (Thermo Fisher), and 200 U of RNase Inhibitor (MoloX, Berlin)]. The lysate was transferred to a 2 ml FastPrep tube with Lysis Matrix E (MP Bio) and lysed using a FastPrep-24 instrument for 15 seconds at 4 m/s. The lysate was clarified by centrifugation at 4 °C and 16,100 x g for 10 min. Next, 12 A_260_ units of the lysate were digested with 20,000 U of MNase (New England Biolabs) in the lysis buffer supplemented with 10 mM CaCl_2_ and 500 U RNase Inhibitor. Polysome digestion was performed at 25°C with shaking at 1,450 rpm in a heating block for 90 min (MNase +). As a control, 12 A_260_ units of undigested lysate were also included to confirm the presence of intact polysomes (MNase -). The MNase digestion was stopped using ethylene glycol-bis(β-aminoethyl ether)-N, N, N′, N′-tetraacetic acid (EGTA) at a final concentration of 6 mM.

To analyze polysome profiles, 12 A_260_ units were layered onto a linear 10%-55% sucrose gradient prepared in the following buffer: [10 mM MgCl_2_, 20 mM Tris-HCl, pH 7.5, 100 mM NH_4_Cl, 5 mM CaCl_2_, 1 mM dithiothreitol (DTT)], in an 13.2 ml open-top polyclear tube (Seton Scientific). Gradients were centrifuged at 35,000 rpm for 2 h and 30 min at 4°C in a Beckman Coulter Optima XPN-80 ultracentrifuge using a SW40-Ti rotor. The gradients were processed using a Gradient Station (IP, Biocomp Instruments) fractionation system with continuous absorbance monitoring at 254 nm to resolve the free RNA, ribosomal subunit, and polysomes peaks. Simultaneously, a UV profile at 254 nm was recorded.

The 70S monosome fractions were manually collected and subjected to RNA extraction to purify the RNA footprints. The RNA was extracted from the fractions or cell pellets for total RNA using hot phenol– chloroform–isoamyl alcohol (25:24:1, Roth) or hot phenol (Roth), respectively, as previously described (51, 52). Ribosomal RNA was depleted from 1 µg of DNase I-digested total RNA by using hybridization probes (Lexogen, Germany) according to the manufacturer’s protocol and fragmented (Ambion 10 x Fragmentation Reagent). Monosome footprints and fragmented total RNA were size-selected (26–34 nt) on a 15% polyacrylamide/7 M urea gel as described previously (24) using RNA oligonucleotides NI-19 and NI-20 as references. RNA was cleaned up and concentrated by isopropanol precipitation with 15 μg GlycoBlue (Ambion) and dissolved in water. Libraries were prepared by the Core Unit SysMed (University of Würzburg) using an adapter ligation protocol without fragmentation and were sequenced on a NextSeq 2000 instrument (high output, 100 cycles) at the Core Unit SysMed at the University of Würzburg.

### General processing of ribosome profiling data

Sequencing data were processed and analyzed using HRIBO (version 1.8.1) (53). The reference genome of *Escherichia coli* CFT073 (ASM744v1) and its corresponding annotation file (GCA_000007445.1) were obtained from NCBI. The annotation of sRNAs in this strain was obtained from a previous publication reannotating the CFT073 genome (54). Adapter sequences and low-quality fragments were trimmed with Cutadapt v. 1.9 (55) from sequencing libraries before mapping reads to the reference genome. Residual rRNA reads following depletion, as well as reads mapping to tRNA, were computationally removed with Samtools (56). Computational removal of rRNA and tRNA reads was performed to eliminate highly abundant background signals, thus reducing noise and enhancing the accuracy of translation-specific read identification and analysis.

Subsequent analysis included quality control, coverage analysis of annotated genomic features, and ORF prediction, all of which was done within HRIBO (v1.8.1). Translational efficiency (TE) was evaluated in HRIBO using a customized annotation set comprising CDS, annotated sORFs, sRNAs, tRNAs, and rRNAs. Specifically, TE was calculated as the ratio of median-of-ratios normalized Ribo-Seq counts to RNA-Seq counts. Only features achieving reads per kilobase of transcript per million mapped reads (RPKM) of at least 10 were included to ensure robustness of translational efficiency estimates.

### ORF predictions and filtering

ORF predictions were performed using two tools integrated in HRIBO (v1.8.1): DeepRibo (v1.1) (57) and REPARATION_blast (v1.0.9) (58), a BLAST-based adaptation of REPARATION. Results from both tools across replicates and conditions were compiled and annotated with metrics such as expression levels, translational efficiency (TE), and DeepRibo scores. Predicted ORFs with 100 codons or fewer were considered as sORFs. Predicted ORFs with no overlaps with annotated coding regions were considered as intergenic ORFs. Candidate sORFs were selected by applying cutoff criteria, requiring a mean TE ≥ 1, RPKM ≥ 10, and DeepRibo score ≥ 0. The resulting intergenic sORFs were analyzed using TMHMM (v2.0) (59) to predict transmembrane helix topology. The final set of sORFs was manually curated based on coverage profiles and predicted topology. The raw data from Ribo-seq and RNA-seq have been deposited as described in the **Data availability** section.

### Whole cell proteome analysis

The proteomics procedure was applied as described in detail previously (60). Cell pellets were resuspended in 300 μl lysis buffer (2% sodium lauroyl sarcosinate (SLS), 100 mM ammonium bicarbonate) and heated for 10 min at 90°C. The amount of proteins was determined by bicinchoninic acid protein assay (Thermo Scientific). Aliquots of total lysate were further processed to enrich the small protein fraction as described previously with modifications (61). Briefly, aliquots containing 100 µg of total protein lysate in lysis buffer (100 mM triethylammonium bicarbonate (TEAB), pH 8.0, supplemented with an EDTA-free protease inhibitor tablet) were concentrated to dryness by vacuum evaporation at 45 °C. The dried samples were reconstituted in 20 µL of 420 mM TEAB, followed by the slow addition of 40 µL acetonitrile containing 0.1% trifluoroacetic acid (TFA). Samples were vortexed for 30 s and incubated at 20 °C with shaking at 1,300 rpm for 1 h. After centrifugation at 21,000 × g for 20 min at 20 °C, the supernatant was collected and dried by vacuum evaporation.

Proteins from total cell lysate and small protein enriched lysate were reduced with 5 mM Tris(2- carboxyethyl) phosphine (Thermo Fisher Scientific) at 90°C for 15 min and alkylated using 10 mM iodoacetamid (Sigma Aldrich) at 25°C for 30 min in the dark. For tryptic digestion, 50 µg protein was incubated in 0.5% SLS and 1 µg of trypsin (Serva) at 30°C overnight. Post digest, detergent was precipitated by acidification and peptides were purified using C18 solid-phase extraction cartridges (Macherey-Nagel).

Peptides were analyzed using liquid chromatography followed by mass spectrometry and carried out on an Exploris 480 instrument connected to an Ultimate 3000 RSLC nano and a nanospray flex ion source (all Thermo Scientific). The following separating gradient was used: 94% solvent A (0.15% formic acid) and 6% solvent B (99.85% acetonitrile, 0.15% formic acid) to 25% solvent B over 40 minutes, and an additional increase of solvent B to 35% for 20min at a flow rate of 300 nl/min.

MS raw data was acquired on an Exploris 480 (Thermo Scientific) in data independent acquisition (DIA) mode. The funnel RF level was set to 40. For DIA experiments full MS resolution were set to 120.000 at m/z 200. AGC target value for fragment spectra was set at 3000%. 45 windows of 14 Da were used with an overlap of 1 Da between m/z 320-950. Resolution was set to 15,000 and IT to 22 ms. Stepped HCD collision energy of 25, 27.5, 30 % was used. MS1 data was acquired in profile, MS2 DIA data in centroid mode.

Analysis of DIA data was performed using the DIA-NN version 1.8 (62) using a uniprot protein database from *E.coli* with additionally added small protein sequences to generate a data set specific spectral library for the DIA analysis. The neural network based DIA-NN suite performed noise interference correction (mass correction, RT (retention time) prediction and precursor/fragment co-elution correlation) and peptide precursor signal extraction of the DIA-NN raw data. The following parameters were used: Full tryptic digest was allowed with two missed cleavage sites, and oxidized methionines and carbamidomethylated cysteines. Match between runs and remove likely interferences were enabled.

The precursor FDR was set to 1%. The neural network classifier was set to the single-pass mode, and protein inference was based on genes. Quantification strategy was set to any LC (high accuracy). Cross-run normalization was set to RT-dependent. Library generation was set to smart profiling. DIA-NN outputs were further evaluated using the SafeQuant (63, 64) script modified to process DIA-NN outputs. The mass spectrometry proteomics data have been deposited to the ProteomeXchange Consortium via the PRIDE partner repository with the dataset identifier PXD065547. The proteomics data have been deposited as described in the **Data availability** section. (65).

### Cell-free synthesis

The experiments were performed as described previously (43, 44). Briefly, the PURExpress kit (NEB) was used to mimic the transcription and translation reaction *in vivo*. Lipid sponge droplets were included in the cell-free system, providing a hydrophobic environment with a large membranous surface area. For this purpose, a uniform lipid film was first formed by evaporating a mixture of chloroform and 10 mM N-oleoyl b-D-galactopyranosylamine (GOA, synthesized by GLYCON Biochemicals) under nitrogen gas. The lipid film was then rehydrated by vortexing with a rehydration solution containing 90 mM octylphenoxypolyethoxyethanol (IGEPAL, Sigma) to form lipid sponge droplets. To initiate protein synthesis, 0.5 µl of a DNA template solution, achieving a final concentration of 9.5 nM, was added to other cell-free components together with lipid sponge droplets. The reactions were carried out for 3 hours at 37 °C.

### Fluorescence microscopy

For the *in vitro* experiments, a 4.5 µl aliquot of the cell-free reaction mixture was pipetted into a Lumox dish (Sarstedt) and sealed with a cover glass. The reaction was incubated at 37 °C on a microscope stage and observed using a Nikon Eclipse Ti-E inverted fluorescence microscope with a 40× objective lens. The images were captured after 3 hours of incubation and analyzed with NIS-Elements Viewer. The propensity of the synthesized small protein towards a hydrophobic environment was quantified by the ratio of green fluorescence inside to outside the lipid sponge droplets. For the *in vivo* expression experiments, *E. coli* K-12 MG1655 carrying plasmids pLW73–pLW83 was grown overnight in LB supplemented with ampicillin, diluted 1:100 into fresh medium, and grown to OD₆₀₀ ≈ 0.6. Cells were washed with the tethering buffer and immobilized on 1% agarose pads prepared in the same buffer. mNeonGreen fluorescence was imaged using a Nikon Ti-E inverted fluorescence microscope equipped with a 100× oil-immersion objective, SOLA nIR LED light source, Orca Fusion sCMOS camera, perfect focus system, and NIS-Elements AR software.

### Flow cytometry

Cells were grown as described above and exposed at OD₆₀₀ ≈ 0.6 to the following stress conditions: 500 mM NaCl (high salt), 5 mM H₂O₂ (oxidative stress), or 0.05% SDS (membrane perturbation). Heat stress was applied from the start of the culture. After 2 h of stress exposure, cells were diluted in tethering buffer, and mNeonGreen fluorescence was measured using a BD LSR Fortessa SORP flow cytometer with 488-nm excitation. Data were analyzed as described previously (66).

### Genomic neighborhood and conservation analysis

Candidate sORFs were mapped from the CFT073 reference genome (AE014075.1) to the genome panel with BLASTN from BLAST+(67) (E-value ≤1e-3; minimum identity 60%; blastn-short for queries <50 bp). GenBank annotations were parsed with Biopython (68), and the five annotated CDS features upstream and downstream of each mapped sORF were extracted. Neighboring proteins were assigned to canonical orthogroups with OrthoFinder (67). Conserved genomic neighborhood scoring (CGNS) was computed against either a consensus template, defined by the most frequent orthogroup at each relative CDS position among scoring-included genomes, or against the CFT073 reference-neighborhood template. In both cases, species-level CGNS combined exact positional presence, relative order conservation, and nearest-position distance within the local window, following prior quantitative synteny and gene-neighborhood scoring approaches (69–73).

Mapped sORF hits were extended in frame with Biopython sequence-translation routines to the first terminal stop codon (TAA, TAG, or TGA). Start sites were selected by a peptide-guided local scan: all ATG, GTG, and TTG starts within 20 bp of the BLASTN hit were translated, initiator-normalized to methionine, and ranked by longest N-terminal match to the CFT073 peptide, then by overall identity and absence of premature stops; nucleotide-frame inference was used when peptide matching was uninformative. Non-reference hits were excluded from CGNS only when ORF extension failed or when the inferred ORF was shorter than two-thirds of the CFT073 ORF. Longer non-reference ORFs, terminal stop codons, opposite-strand CDS overlaps, and possible CFT073 truncations were retained as annotations. Ribo-seq support was assessed by merging CGNS/ORF annotations with the original translation efficiency, Ribo-seq RPKM, Reparation, and DeepRibo fields. The formulae used for the analysis are listed in Table S4. The CGNS script has been deposited as described in the **Data availability** section. Neighborhood maps were generated from GenBank-derived CDS coordinates, with directional gene arrows positioned relative to each sORF and colored by OrthoFinder orthogroup.

The TBLASTN search was performed by submitting individual amino acid sequences against NCBI core_nt through the NCBI website (E-value ≤ 0.05, BLOSUM62, gap costs 11/1, SEG filtering, composition-based statistics mode 2, and returned subjects ≤ 5,000). The returned sequences were subject to start/stop and translated protein sequence analysis. All high-scoring segment pairs (HSPs) were counted separately using strand-oriented nucleotide sequences retrieved with NCBI EFetch, classifying the first aligned codon as ATG, GTG, TTG or other and the codon immediately following the HSP as TAA, TAG, TGA or other. These measurements describe alignment boundaries rather than experimentally established ORF boundaries. To generate the sequence logos, one HSP per returned subject was selected by lowest E- value per genome, with ties resolved by highest bit score, greatest query coverage and lowest HSP number. Subject residues were projected onto query positions, omitting subject insertions and treating gaps, uncovered positions and X as missing, while retaining translated stops. Letter heights were calculated as symbol frequency multiplied by small-sample-corrected information content over 20 amino acids plus the stop symbol and positional occupancy (the fraction of selected subjects contributing a symbol). Minimum occupancy was the lowest positional occupancy across the query. The exact TBLASTN query sequences and search parameters are available in the GitHub repository (https://github.com/JustABiologist/synteny-smORF/tree/main/data).

## Results

### Ribo-seq and RNA-seq analyses of uropathogenic *E. coli*

To obtain detailed information on the transcriptome and translatome of the UPEC strain CFT073, we applied RNA-seq and Ribo-seq techniques (Fig. 1A). Briefly, we grew the bacterial cells to early log phase and then harvested them by fast-chilling in an ice-water bath to reduce ribosome run-off and capture actively translating ribosomes on mRNA. One set of samples was processed for total RNA isolation and transcriptomic analysis. The other set of samples was lysed and proceeded to ribosome purification. We observed defined peaks of the monosome and polysomes in the gradient centrifugation, indicating that actively translating ribosomes were successfully isolated (Fig. 1B). MNase treatment was optimized to digest RNAs that were not protected by ribosomes and converted the polysome populations to monosomes (Fig. 1B). After the depletion of rRNAs, the extracted RNA samples were subjected to next-generation sequencing.

**Fig. 1.**
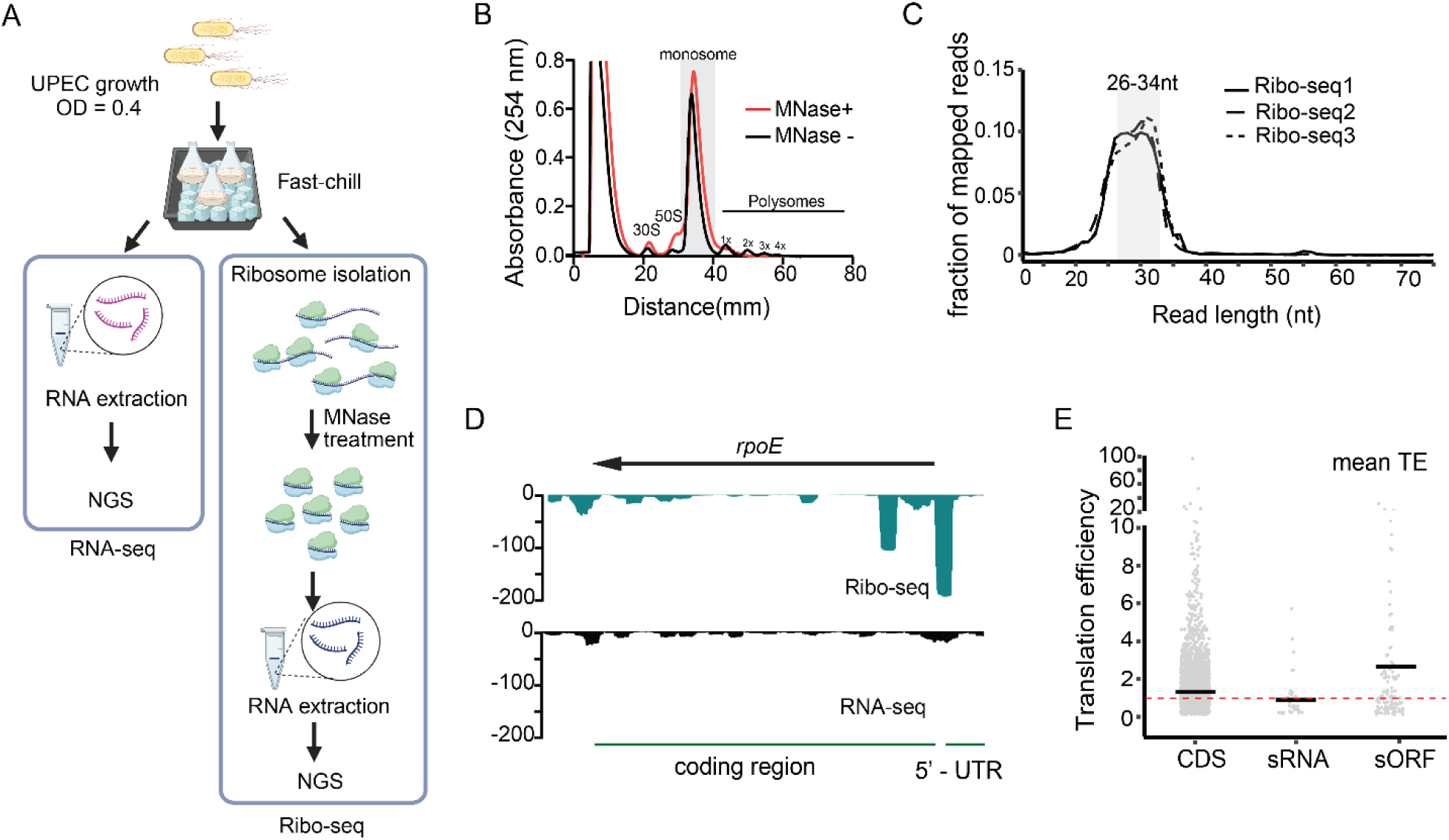
Determination of the UPEC transcriptome and translatome by RNA-seq and Ribo-seq, respectively. (**A**) The experimental workflow started with the growth of wild-type UPEC CFT073 followed by halting translating ribosomes via fast chilling. Samples were divided into two sets. One set was used for RNA extraction followed by cDNA preparation and next-generation sequencing (NGS). The other set of cells were lysed and ribosomes were isolated. Polysomes were digested into monosomes with MNase (MNase +). The monosome was then used for total RNA extraction. The isolated RNA was then subjected to cDNA library preparation and NGS. (**B**) Quality control during the Ribo-seq experiment. Undigested (MNase -) and MNase-treated (MNase +) fractions were analyzed to confirm the presence of stable polysomes before digestion, the digestion of the polysomes into monosome fraction, and the enrichment of the monosome fraction after digestion. (**C**) Read length distribution for libraries from three replicates of the Ribo-seq experiment. The size-selected lengths (26-34 nt) are highlighted with grey boxes. (**D**) JBrowse genome browser shows that by comparing Ribo-seq and RNA-seq, the coding region and 5’UTR of a gene (i.e. *rpoE* [2,972,304..2,972,912 (-)] can be defined based on ribosome footprints. (**E**) Translation efficiency (TE, i.e., the fraction of normalized Ribo-seq counts divided by normalized RNA-seq counts, see methods) for different feature classes in the *E. coli* CFT073 annotation. A total RNA RPKM of ≥10 was required. CDS: coding DNA sequences. sRNA: small RNA. sORF: small open reading frame. Panel A was created using BioRender. *Schematic workflow and experimental data of Ribo-seq coupled with RNA-seq*

We used the reference genome of CFT073 from NCBI and non-coding RNA sequences from a previous reannotation to analyze the sequencing data (54). The latest version of HRIBO (version 1.8.1) was the main workhorse for data processing, with the workflow published previously (53). HRIBO was developed in 2020 and has been used for analyzing Ribo-seq data obtained from diverse prokaryotes, including *Salmonella enterica* Typhimurium, *Sinorhizobium meliloti*, *Methanosarcina mazei*, and *Campylobacter jejuni* (6, 9, 10, 50). Inspection of the raw reads revealed the successful removal of adapters, rRNA, and tRNA contaminations, indicating high quality of the samples and sequencing data. The majority of reads had the length of 26-34 nucleotides, consistent with appropriate nuclease digestion and the size selection (Fig. 1C). When comparing the resulting Ribo-seq read coverage data to paired RNA-seq data, we obtained detailed information regarding a specific ORF, such as ORF boundaries, 5’-UTR, 3’-UTR and translational efficiency (TE: Ribo-seq/RNA-seq) (Fig. 1D). Inspection of protein-coding ORFs and known non-coding RNAs further confirmed the success of the Ribo-seq and RNA-seq data. On average, the translational efficiency of sORFs and annotated protein-coding DNA sequences (CDS) was greater than 1, while that of sRNAs was less than 1 (Fig. 1E). Therefore, we used TE ≥ 1 as one of the criteria for sORF prediction, together with a minimum RNA RPKM threshold to ensure reliable TE estimation.

### Exploring translation of annotated genes

UPEC CFT073 has a total of 5490 annotated ORFs in the genome. Using mass spectrometry, we detected 2115 annotated proteins when UPEC cells were grown under the same conditions and harvested in log phase. The RNA-seq data detected transcripts from 3411 annotated ORFs with RPKM set at 10 or higher. Of these, 2706 transcripts had ribosomal footprints indicating translation with a TE value above 1 (Fig. 2A). The number of translated ORFs (2706) was slightly higher than the 2115 proteins detected by the conventional proteomic approach, indicating an enhanced sensitivity of Ribo-seq. With the Ribo-seq raw data as input, DeepRibo, a neural network-based prokaryotic annotation tool, detected a total of 2130 annotated ORFs, of which 1933 ORFs had a positive confidence score (DeepRibo ≥ 0). Of them, 83.4% (1611 out of 1933 ORFs) passed the other filtering criteria (RPKM ≥ 10, TE ≥1). These results indicated that DeepRibo ≥ 0 is a stringent filtering condition. With this filtering condition, the DeepRibo prediction was slightly more conservative but largely consistent with Ribo-seq and proteomics data for annotated proteins (Fig. 2A).

**Fig. 2.**
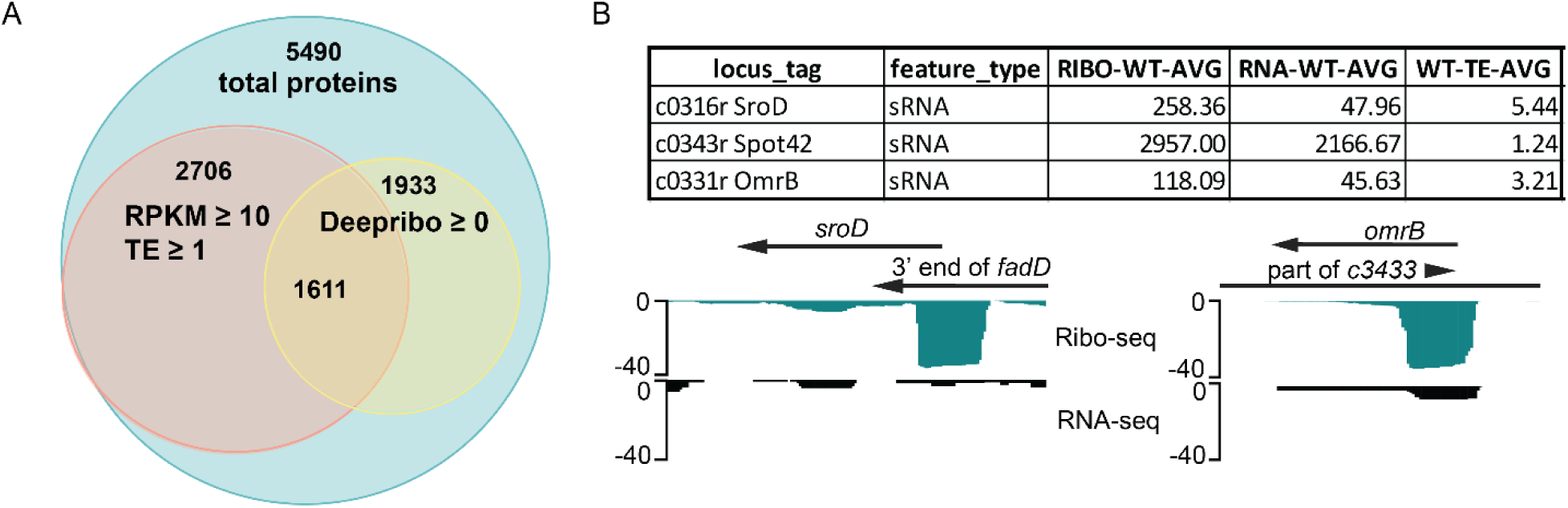
The translatome of annotated genes. (**A**) Comparison of filtering criteria. With the cutoff of RPKM ≥ 10 and TE ≥ 1, Ribo-seq coupled with RNA-seq identified 2706 of the total 5490 annotated genes in the genome. Using the same input data set, with DeepRibo ≥ 0, 1933 annotated genes were identified. (**B**) Three small RNAs, Spot42, SroD, OmrB showed TE values above 1 as shown in the table. The representative Ribo-seq and RNA-seq profiles of *sroD* [2,039,672..2,039,791(-)] overlaps with the 3’ end of *fadD*. The *omrB* gene [3,284,337..3,284,418 (-)], which encodes the OmrB sRNA, is located on the non-template DNA strand within the c3433 gene. *Schematic and experimental data showing the identified translatome and protein-coding sRNAs*

UPEC CFT 073 has a total of 46 annotated sRNAs. Most of them showed no or low Ribo-seq reads with TE < 1, as expected. One of the exceptions was Spot 42, which was previously identified as a dual-function sRNA in the *E. coli* K-12 strain (74). Spot 42 acts as a small RNA and encodes the small protein SpfP (15 aa). SpfP binds to the global transcription regulator CRP (cyclic adenosine monophosphate receptor protein) and represses specific promoters, reinforcing catabolic repression (74). Interestingly, two sRNAs (SroD and OmrB) showed notably high translation efficiency with TE scores ranging from 3 to 5 under the tested growth condition (Fig. 2B). They had much lower normalized reads in Ribo-seq and RNA-seq when compared to those of Spot 42, but well above the cutoff of 10. SroD has an unknown function so far. In the K-12 strain, it was found to be expressed in a narrow window when cells entered the stationary growth phase (75). The small RNA OmrB interacts with Hfq and is known to be involved in the modulation of outer membrane protein production at the posttranscriptional level in *E. coli* and *Salmonella* (76). Closer inspection of the UPEC CFT073 genome and the sequencing data revealed that the *sroD* gene overlaps with the 3’ end of the *fadD* gene (Fig.2B). Therefore, the Ribo-seq reads from *sroD* and *fadD* could not be differentiated from each other. The *omrB* gene, which encodes an 82 nt RNA, is located on the non-template DNA strand within *c3433*, a gene for a putative membrane protein, indicating that the high TE score results from the translation from *omrB*.

### Identification of unannotated putative small proteins

We used the NCBI reference genome and a published reannotation of the UPEC strain CFT073 as inputs, together with a database of all possible ORFs encoded in the UPEC genome. HRIBO, integrating DeepRibo and REPARATION_BLAST deep-learning models, was then applied to predict translation initiation sites and ORFs based on DNA sequence features, ribosome profiling density, and homology information, assigning a probability score (DeepRibo score) to each candidate ORF. The whole genome survey identified 561 sORFs (less than 100 codons) after filtering with the stringent criteria (RPKM ≥ 10, TE ≥ 1, and DeepRibo ≥ 0). Of 561 identified sORFs, 410 were not annotated so far, 138 of which were located in intergenic regions of the genome. To have an overview of the Ribo-seq reads, we performed metagene profiling and mapped the 5’ and the 3’ ends of the reads (Fig. S1). Major peaks were observed at positions near −22 and +8 relative to the start codon in the 5′ and 3′ read mappings, respectively (Fig. S1A), indicating a 7-nucleotide shift compared with the canonical ribosomal occupancy pattern (−15 and +15). We validated our algorithm using additional published *E. coli* ribosome profiling datasets (Fig. S1B, C). This shift persisted after excluding highly expressed genes and across different read-lengths, indicating that it was not driven by specific transcripts or incomplete nuclease digestion. It may instead reflect over-digestion, effects of footprint length selection, library preparation, other sample processing steps, and/or underlying biological factors. Adding this information to our prediction pipeline, the second round analysis resulted ORFs about 85% identical to the list generated from the first round (Fig. S2). We then focused on the 477 putative sORFs identified by both analyses, of which 338 were unannotated and 113 were located in intergenic regions.

The length distribution of the unannotated sORFs differs markedly from that of the annotated ones (Fig. 3A). The former group had the most sORFs between 50-75 codons in length. In contrast, the majority of annotated sORFs were longer, with codon numbers ranging from 75 to 100. Consequently, the unannotated group contained a higher percentage of shorter sORFs (10-50 codons) than the annotated group (29% vs. 6%). Next, we analyzed the codon usage of the unannotated sORFs and compared it to that of the annotated group (Fig. 3BC). Both groups had ATG as the predominant start codon (79% and 90%, respectively). However, the unannotated sORF group had a slightly higher percentage of the GTG start codon (15% vs. 6%). Regarding the stop codons preference, the annotated group favored TAA over TGA (60% vs. 44%), while the unannotated group used the two stop codons almost equally (TAA: 44%, TGA: 41%) (Fig. 3C). We did not observe a significant difference between these two groups regarding translation efficiency (Fig. 3D).

**Fig. 3.**
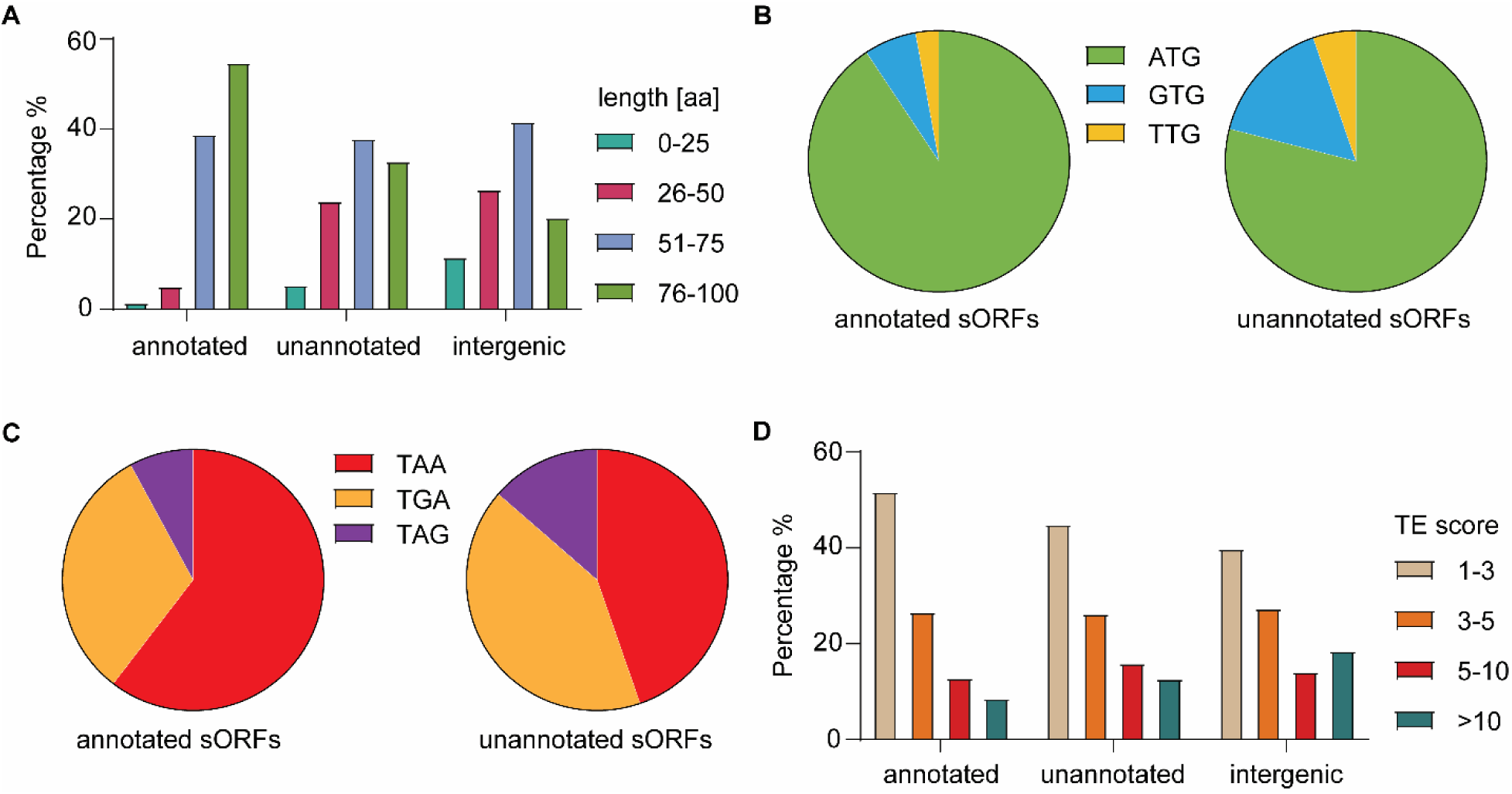
Analyses of identified sORFs. Three groups of sORFs, the annotated, the unannotated and the intergenic sORFs were compared regarding their length distribution (**A**), codon usage (**B** and **C**), and TE distribution (**D**). *Analytical data showing the length distribution, start and stop codon usage and translation efficiency of identified sORFs*

We then focused on the 113 sORFs located in intergenic regions of the genome, as our Ribo-seq approach could not distinguish translation from a putative sORF, if it was nested in or overlapped with other coding DNA. Overall, the intergenic subset showed a slightly higher translation efficiency when compared with the total unannotated sORFs (Fig. 3D). This mainly resulted from the slight increase in the percentage of sORFs with TE ≥ 10. This higher average TE value (6.8 vs. 5.4) suggested that intergenic sORFs are likely translated more robustly. The intergenic subset shared similar start/stop codon preferences with the total unannotated sORFs. The AUG was predominantly used as the start codon (81%), and TGA and TAA were used almost equally as the stop codons (42% and 43%, respectively). Regarding the translated small protein length, intergenic sORFs encode more proteins in the length of 25 – 75 aa when compared to those translated from the total unannotated sORFs (70% vs. 64%).

We manually inspected the 113 intergenic sORFs, excluded the ones that showed TE > 1 only due to less than 30% of the entire ORF, and the ones with RNAseq reads lower than 10 in each replicate. For the resulting 78 sORFs with higher confidence, we then searched for their homologs using the NCBI BLAST tool with the protein sequences as query (77). We found 15 of them annotated as known proteins in the genome of other microorganisms. The remaining intergenic sORFs encoded uncharacterized, hypothetical, and novel small proteins.

### Validation of putative small proteins by mass spectrometry

To validate translation of the putative small open reading frames (sORFs) at the protein level, we performed mass spectrometry–based proteomic analyses. Protein sequences corresponding to all putative sORFs were added to the *E. coli* CFT073 proteomic database downloaded from NCBI to enable their identification. Cell lysates were prepared using two complementary approaches. In addition to a conventional detergent-based protocol for generating total cell lysates from uropathogenic *E. coli* cells, we employed a size-enrichment strategy to improve the detection of small proteins. Specifically, larger proteins were selectively precipitated using acetonitrile, thereby enriching the small-protein fraction in the supernatant (see Materials and Methods for details). The resulting supernatants were concentrated and subjected to liquid chromatography–tandem mass spectrometry analysis.

Using these approaches, peptide fragments corresponding to five small proteins were consistently detected in both rounds of mass spectrometry analysis, supporting their expression under the tested growth conditions. In addition, two small proteins were detected in only one of the two preparation methods (File S1). Notably, none of the validated small proteins were predicted to contain transmembrane helices, indicating that small membrane proteins remain particularly challenging to detect by mass spectrometry–based proteomic workflows.

### Identification of putative small membrane proteins using a cell-free system

Many bacterial small proteins that have been functionally characterized are membrane-associated small proteins, containing a single transmembrane helix (5, 12). We used the TMHMM (v2.0) algorithm (59) to predict the presence of transmembrane helices and identified 25 small membrane proteins in the annotated group, including the well-characterized small protein MgrB (also named as YobG). With the same approach, we inspected the 78 intergenic encoded small proteins and identified 15 of them that had at least one putative transmembrane helix (File S2). They included the recently characterized small membrane protein MgtS, YohP, and homologs of type I toxins (35, 36, 78, 79). Of the 15 putative small membrane proteins, seven have known or expected functions and two are annotated but not characterized. The remaining six have not been previously identified or annotated in genomes of other organisms. Interestingly, four out of the six unknown proteins (SMP3, SMP7, SMP11, and SMP14) have their coding genes residing within the PAIs of the UPEC genome (predicted by IslandViewer 4) (80).

To confirm their membrane incorporation, we expressed all six putative novel small proteins (SMP3, SMP7, SMP10, SMP11, SMP12, and SMP14) in parallel with six small membrane proteins annotated in other *E. coli* strains (YfgG, MgtS, PmrR, Hok/Gef, Blr, and HokA) in a cell-free system containing lipid sponge droplets. This cell-free system was shown to produce functional small membrane proteins (48). The expression could be easily visualized under a microscope when a small protein was fused with the fluorescent protein mNeonGreen (48). Due to the hydrophobic nature of transmembrane helices, newly synthesized membrane proteins partitioned from the aqueous reaction mixture into lipid sponge droplets, leading to higher green fluorescence within the droplets than in the surrounding solution.(48, 81). We constructed sORF-*mneongreen* fusion templates and followed green fluorescence during cell-free synthesis (Fig. 4). The synthesis of the mNeonGreen-tagged known small membrane protein, SafA, was used as a positive control, while mNeonGreen alone and mNeonGreen tagged small cytosolic ribosomal proteins (RpmG and RpmH) as negative controls.

**Fig. 4.**
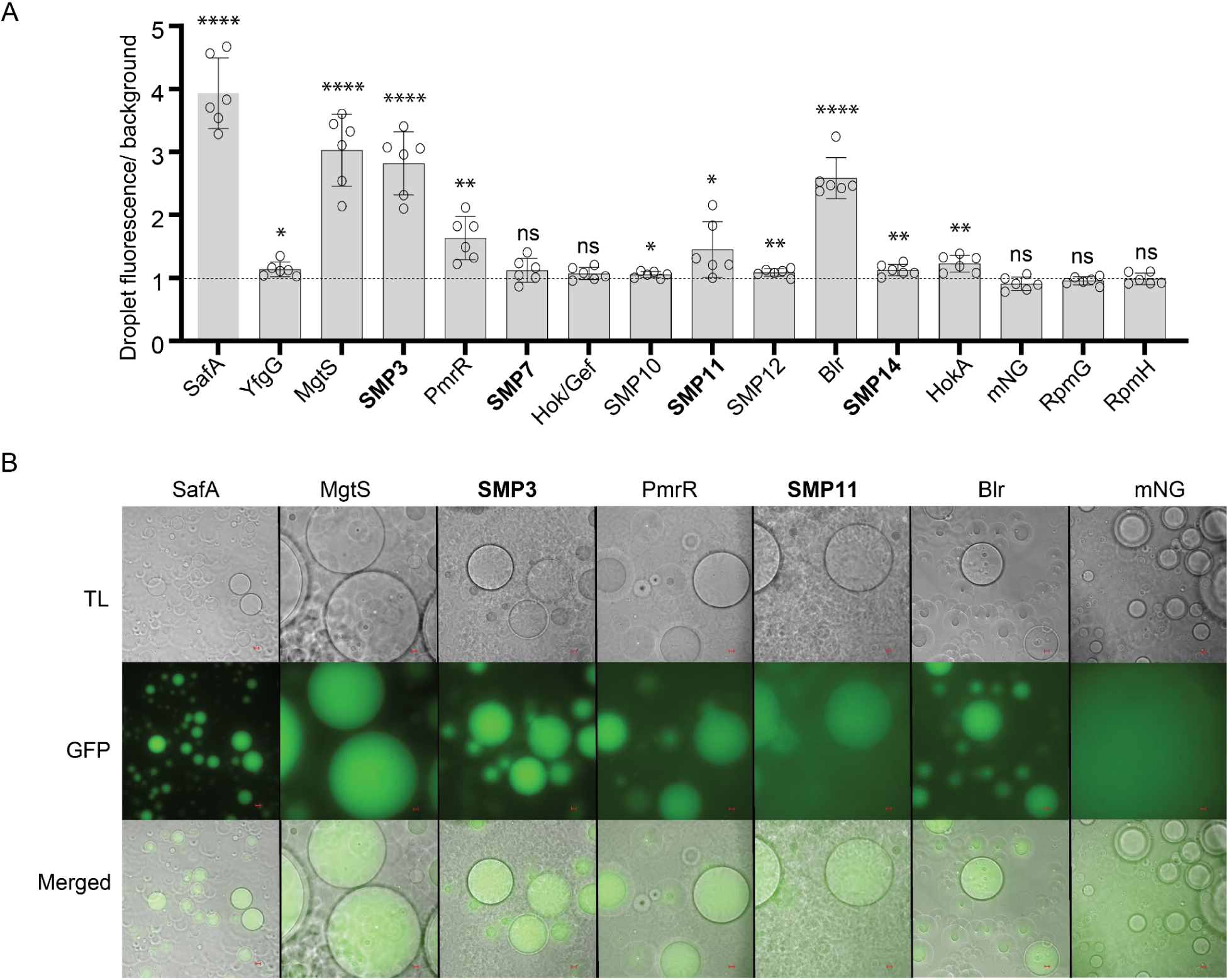
Cell-free synthesis of predicted small membrane proteins (SMP). (**A**) The mNeonGreen-labeled SMPs were produced in the cell-free system containing lipid sponge droplets. The propensity towards a hydrophobic environment was quantified by the ratio of fluorescence inside to outside the droplets. SMPs in bold font have their genes residing within PAIs. The significance of fluorescence change (inside vs. outside droplets) was analyzed using an unpaired t-test. (**B**) Representative images from the transmitted light (TL), GFP fluorescence channel, and the merged channels are shown (scale bar represents 10 µm). Data are representative of at least two independent experiments. mNG, mNeonGreen; ****, P < 0.0001; ***, P < 0.001; **, P < 0.01; *, P < 0.05; ns, not significant. *Experimental data showing detectable protein synthesis and membrane localization of MgtS, SMP3, PmrR, SMP11, Blr in a cell-free system with lipid sponge droplets*.

The positive control showed strong green fluorescence localized in the lipid droplets after three hours of incubation as expected. The fluorescence inside the droplets was approximately 4-fold higher than that outside the droplets. All three negative controls revealed no significant difference in green fluorescence between the inside and outside of the lipid sponge droplets. Three characterized small membrane proteins (MgtS, PmrR, Blr) showed significantly higher fluorescence inside the droplets, consistent with their known localization in vivo. YfgG, an uncharacterized small protein involved in nickel and cobalt stress, showed only slight preference for the hydrophobic droplets, likely due to a charge residue (Arg) within its predicted transmembrane region. HokA and Hok/Gef family proteins are known for oligomerizing and pore forming within the bacterial inner membrane. They showed only slight or no preference to localize to the droplets that have an extensively convoluted membranous surface instead of a planar bilayer in a cell.

Of the six putative small membrane proteins, SMP3 and SMP11 showed a relatively stronger tendency to localize in lipid sponge droplets with the fluorescence ratios ranging from 1.5 to 3, while SMP7 did not show significant difference in green fluorescence between the inside and outside of the lipid sponge droplets. SMP10, SMP11, SMP12 and SMP14 demonstrated a slight but significantly higher fluorescence

inside the hydrophobic droplets, suggesting a small likelihood of membrane localization (Fig. 4 and Fig. S3). SMP3, SMP7, SMP11, and SMP14 (Fig. 4A in bold) were particularly interesting due to the genomic location of their coding genes within pathogenicity islands.

### Validation of putative small membrane proteins *in vivo*

To validate the expression of the six putative small membrane proteins (SMP3, SMP7, SMP10, SMP11, SMP12, and SMP14) *in vivo*, we cloned the predicted coding sequence together with about 200 bp upstream UTR and promoter region into a high-copy plasmid and fused mNeonGreen to the C terminus of the protein. A plasmid with the gene of mNeonGreen alone without any promoter sequence and sORF served as a negative control, while a plasmid with the known small membrane protein Blr was used as the positive control. We also included a construct with the previously detected but not characterized small membrane protein YohO (29). *E. coli* cells with the constructed plasmids were grown to exponential phase and examined for the presence of mNeonGreen with fluorescence microscopy. The level of fluorescence was further quantified with flow cytometry. As expected, the negative control showed very faint fluorescence (Fig. 5A). The positive control Blr showed strong fluorescence at the cell periphery, but not at the septum in the presence of the wild-type genomic copy, consistent with the previous report (82). Cells carrying the *smp7-* and *yohO-*containing plasmids showed strong fluorescence throughout the entire cell volume and at cell periphery, respectively, confirming *in vivo* expression of these small proteins with distinct localization patterns (Fig. 5A). SMP7 did not localize to lipid sponge droplets in the cell-free system (Fig. 4A), consistent with these *in vivo* results, indicating that SMP7 might be a small cytosolic protein. The previously uncharacterized protein YohO localized to the membrane as a mNeonGreen fusion, consistent with the prediction. No clear membrane localization of SMP3 was detected *in vivo*, which disagrees with the *in vitro* data. This may result from inefficient folding and chromophore maturation of mNeonGreen when in the periplasm, or impairment of SMP3 membrane insertion *in vivo* due to the mNeonGreen tag.

**Fig. 5.**
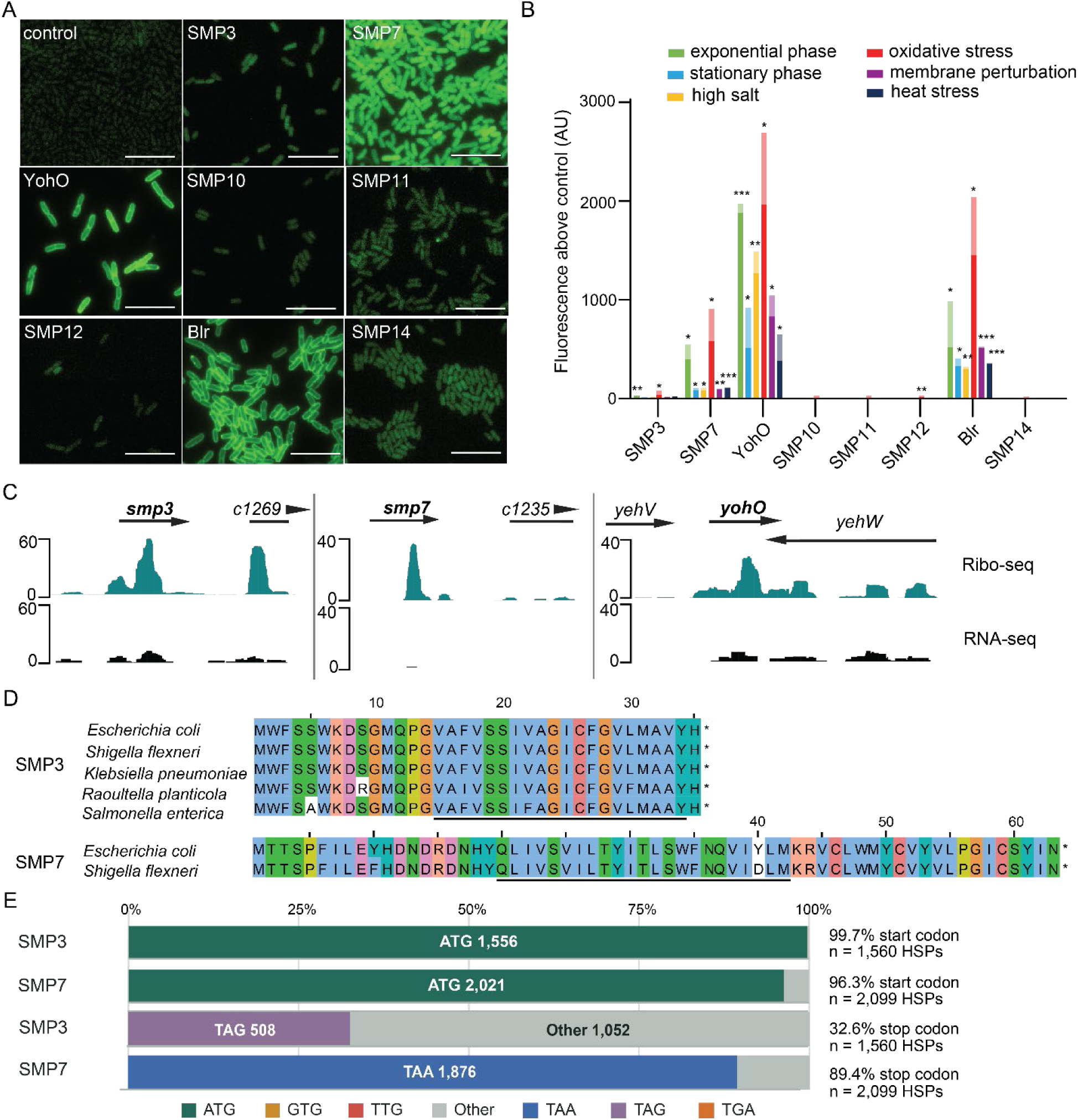
Validation of putative small membrane proteins in vivo. Small proteins were expressed as C-terminal mNeonGreen fusions from their native promoters cloned on a high-copy plasmid and analyzed in *E. coli* K-12 MG1655. (**A**) Fluorescence microscopy images showing expression and localization of mNeonGreen-tagged small proteins in logarithmically growing cells. A plasmid lacking both promoter and sORF was used as a negative control. Scale bars represent 10 micrometers. Images are representative of two independent experiments. (**B**) Flow cytometry–based quantification of small protein expression under the indicated stress conditions. Stress treatments were applied during logarithmic growth, except for heat stress, which was applied from the start of the culture. Fluorescence values of the control cells were subtracted to remove background fluorescence. Bars with different shading denote two independent biological replicates. (**C**) Representative Ribo-seq and RNA-seq profiles of SMP3 [1,219,521..1,219,628(+)], SMP7 [1,184,881..1,185,072(+)], and YohO [2,507,364..2,507,471(+)] are shown using the JBrowse genome browser. (**D**) The DNA sequences of SMP3 and SMP7 were found in other *E. coli* strains and indicated species. Alignments of potential protein homologs are shown with conserved residues marked in different colors using Jalview (83, 84). The underlined region presents the predicted transmembrane helix. (**E**) The results of a TBLASTN search using SMP3 and SMP7 as individual query sequences were analyzed. The distribution of codons at the start and the stop positions, the number of total high-scoring segment pairs (HSPs), and the percentage of start and stop codons are shown. *Experimental data showing the expression of SMP3 and SMP7 in vivo and bioinformatic analysis of their distribution and conservation*

To quantify the fluorescence signals over a larger cell population, we grew the strains to exponential phase and quantified the fluorescence with flow cytometry. Furthermore, we quantified cellular fluorescence at stationary phase and after exposure to stress conditions, including high salt (NaCl), oxidative stress (peroxide), membrane perturbation (SDS), and heat stress (42 °C). We observed fluorescence signals above the negative control from cells carrying *smp3*, *smp7*, *smp12*, *yohO*, and *blr*- containing plasmids, confirming their expression *in vivo* (Fig. 5B, C and S4). Moreover, we observed that oxidative stress repeatedly increased cellular fluorescence, indicating elevated production of small proteins under this stress condition. Other stress conditions led to reduced fluorescence, suggesting lower amounts of small proteins due to either decreased expression or increased protein degradation (Fig. 5B and S4).

SMP3 was found as a DUF6404 family protein in other *E. coli* strains. Homologous sequences of SMP3 were found within hypothetical proteins expressed from PAIs in *Shigella flexneri*, *Klebsiella pneumoniae,* and *Raoultella planticola* (Fig. 5D). In *Salmonella enterica*, the sORF was found in an intergenic region.

Sequence homologs of SMP7 were narrowly distributed and found in *E. coli* and *Shigella flexneri*. A tblastn search in the core nucleotide database (NCBI) resulted in more than one thousand hits for SMP3 and SMP7, majority of which showed ATG as the start codon position of the alignment (Fig. 5E). At the stop codon position, many SMP3 hits appeared to have a sense codon. The translated amino acid sequences from the alignments remained to be highly conserved for both small proteins (Fig. S5A).

### Genomic neighborhood analysis of intergenic sORFs

Due to the low abundance and seemingly conditional expression of the putative small proteins, it was challenging to validate all of them using alternative approaches. Therefore, we sought to analyze their distribution based on the assumption that functional sequences have conserved gene neighborhoods across different strains. We searched for homologous DNA sequences of all 78 intergenic UPEC sORFs in the genomes of ten *E. coli* strains (MG1655, HS, H10407, BW25113, E2437A, EDL933, Nissle 1917, UTI89, E2348/69, MP1), *Shigella Flexneri*, *Salmonella enterica* Typhimurium, *Klebsiella pneumoniae*, and *Raoultella sp*. Sequences with 60% and higher identity were extracted followed by reading frame determination (see Materials and Methods). After translation, homologous proteins shorter than two-thirds of the CFT073 reference due to a nonsense mutation were eliminated. To analyze the genomic context, the information of five upstream and five downstream ORFs of the mapped sORF was extracted and compared to that of the CFT073 sORFs. We further excluded the sORFs located on the non-template strand of a known gene, as genomic context analysis would not be informative for the sORFs in this case. In the end, we identified 37 intergenic sORFs (27 ssolORFs and 10 smemORFs) in CFT073 that had homologous sequences and genomic synteny in at least one other strain (File S3 and S4), supporting their likelihood of encoding functional proteins.

A conserved genomic neighborhood for the top predicted sORFs was identified in 13 and 11 of the analyzed genomes, respectively (Fig. S5B). The sORF encoding SSP1 is located downstream of a pseudogene and upstream of *csrA*, which encodes a global post-transcriptional regulator involved in carbohydrate metabolism. The sORF encoding SSP2 is located between *acrD* (encoding an efflux pump permease) and *yffB* (encoding a putative reductase). In MG1655, SSP2 is annotated as an uncharacterized protein that accumulates during stationary phase and in response to heat shock (29). In contrast, homologous genomic neighborhoods of SMP3 and SMP7 were identified in only two other *E. coli* strains (Fig. S5), although homologous sequences were detected in bacterial species outside the genus *Escherichia* (Fig. 5D). This distinction is not unexpected, as conservation of a genomic neighborhood does not necessarily correlate with sequence conservation of the associated gene and *vice versa*. Nevertheless, conserved genomic neighborhoods may provide additional clues to the potential functions of putative small proteins. In UPEC, the sORF encoding SMP3 is located within a PAI, adjacent to *c1269*, which encodes a secreted, surface-associated lipoprotein implicated in bacterial colonization and biofilm formation (44). The sORF encoding SMP7 is located adjacent to *c1234* and *c1235*, which encode a hypothetical protein and a phage-derived protein, respectively.

## Discussion

*E. coli* is a remarkably diverse species. It includes harmless commensal strains such as MG1655 and strains causing infections in the human intestine, such as EHEC as well as extra-intestinal infections, such as UPEC. Besides the core genes shared among them, pathogenic *E. coli* strains have obtained foreign gene elements via horizontal gene transfer, transposons, phages and plasmids (85). The UPEC strain CFT073 contains no plasmids. Its genome has at least seven prophage-like elements and 60 genomic islands (44, 86). We identified 78 sORFs in the intergenic regions of the UPEC genome, including 15 sORFs coding for putative membrane proteins. After careful manual curation, 58 of these were confirmed to be either hypothetical genes or unique to UPEC. The other 20 sORFs have been found mostly in *E. coli* MG1655 and other closely related enterobacteria. Regarding small membrane proteins, 8 out of 15 (53%) have been identified in other *E. coli* strains, a much higher percentage when compared to that of the total intergenic sORFs. Since many characterized bacterial small proteins are membrane proteins, this could explain the higher prediscovery rate. Nevertheless, the high prediscovery rate of small membrane proteins could also imply that membrane-localized small proteins may be easier to generate in intergenic regions and subject to greater selective pressure, thus evolving faster. Consistent with this hypothesis, a recent evolution model suggested that thymine-rich intergenic regions in yeast have great potential to produce transmembrane helix-containing polypeptides for natural selection (87). In the rice genome, *de novo* genes were found to evolve faster than gene duplicates regarding obtaining new features such as hydrophobicity and secondary structures – physiochemical elements of a TM helix. Smaller molecular weight was also found to be associated with a faster pace of evolution (88).

Four of the six novel small proteins unique to UPEC (SMP3, SMP7, SMP11, SMP14) are encoded by sORFs located inside PAIs of the UPEC genome. Some have multiple rare codons such as AUA for isoleucine, one of the hallmarks of island genes. This could explain why some novel sORFs showed very low protein expression in the cell-free system or below the detection level *in vivo*. The sORFs encoding SMP3, SMP7, and SMP11 are located in the large genomic islands PAI-serX and PAI-pheU (PAI II_CFT073_). CFT073 has thirteen large genomic islands longer than 32 kb. They are involved in UPEC colonization in mice and UPEC virulence (89). PAI-pheU contains about one hundred ORFs, including genes for P-fimbriae, type-1 fimbriae and a large number of hypothetical genes (90). This PAI was found to be prevalent among invasive *E. coli* strains (90, 91). PAI-ser is 113 kb in size, including genes involved in siderophore biosynthesis for iron acquisition, which may enhance UPEC fitness and colonization in the urinary tract (86). It is tempting to speculate that the novel hypothetical sORFs identified in this study, such as SMP3 and SMP7, might be involved in UPEC pathogenicity or host colonization.

In this study, we detected only a few small proteins using the mass spectrometry-based proteomic approach despite the effort of small protein enrichment by removing larger proteins via precipitation. This is not entirely unexpected, as the likelihood of detection reduces when protein sizes decrease.Furthermore, the standard proteomic approach relies on trypsin digestion to produce peptides within the detectable length range. Detection of a protein is highly influenced by its specific sequence. In particular, many small membrane proteins contain a long stretch of hydrophobic residues and have very few trypsin digestion sites, thus challenging to detect. Recently, top-down and bottom-up proteomic approaches have been developed for detecting small proteins and used to validate Ribo-seq discovery in bacteria and archaea (92, 93). These modified proteomic approaches together with alternative proteases would likely assist in validating the UPEC small proteome as well.

We employed plasmid-based expression system as an alternative approach to Western blot for detecting small protein production *in vivo*. By tagging with one of the brightest fluorescent protein (mNeonGreen), we were able to simultaneously observe small protein expression and cellular localization across growth phases and stress conditions. This approach is high-throughput and automation compatible, and likely as sensitive as Western blot, since two known small protein controls were well detected. Its main limitation is the possible impact of the fluorescence protein tag on the function and localization of small proteins. Altogether, our study uncovered a list of previously unknown or uncharacterized intergenic sORFs in the UPEC strain CFT073. The *in vivo* validation presents a high-throughput compatible approach, and provides a foundation for studying those small proteins on their potential roles in UPEC infection.

## Data availability

Ribo-seq data generated in this study have been deposited at the NCBI Gene Expression Omnibus (GEO) under the accession GSE301040. Processed data, as well as an interactive genome browser used for this publication is available on the RIBOBASE (http://www.bioinf.uni-freiburg.de/ribobase/). The mass spectrometry proteomics data have been deposited to the ProteomeXchange Consortium via the PRIDE partner repository with the dataset identifier PXD065547. The script for genomic neighborhood analysis has been deposited in Github (https://github.com/JustABiologist/synteny-smORF).

## Acknowledgements

We thank Dr. Hans-George Koch (University of Freiburg) for protocols and discussions, Dr. Shan Jiang (Max Planck Institute for Terrestrial Microbiology) and Dr. Henrike Niederholtmeyer (Technical University of Munich) for the demonstration of the cell-free synthesis and suggestions to improve the system, Jörg Kahnt (Max Planck Institute for Terrestrial Microbiology) for performing sample preparation for mass spectrometry, the CoreUnit SysMed at the University of Würzburg for help with deep sequencing, members of the SPP2002 and the Yuan group for helpful discussions. Schematic figures were created with BioRender.com. AI-assisted editing was used to improve the readability of the manuscript.

## Author contributions

J.Y., E.F. C.M.S., R.B., designed research; W.L., L.W., E.F., R.G., F.C.G., E.M., T.G., and C.C. performed research; R.G., E.F., W.L., L.W., F.C.G., T.G., and J.Y. analyzed data; and W.L., L.W., E.F., R.G., T.G., F.C.G. and J.Y. wrote the paper with comments from the other co-authors.

## Funding and additional information

This work was supported by the Deutsche Forschungsgemeinschaft (DFG) priority program SPP2002 (SPP2002 YU 247/3-1) to J.Y., SPP2002 Z-project 2 on Ribo-seq and bioinformatics: (BA2168/21-2) to R.B. and Sh580/7-2 to C.M.S.. Moreover, R.B was supported by funds from Germany’s Excellence Strategy (CIBSS - EXC-2189 - Project ID 390939984). This work was supported by the Cloud within the German Network for Bioinformatics Infrastructure (de.NBI) and ELIXIR-DE (Forschungszentrum Jülich and W-de.NBI-001, W-de.NBI-004, W-de.NBI-008, W-de.NBI-010, W-de.NBI-013, W-de.NBI-014, W-de.NBI-016, W-de.NBI-022).

