## supplementary information for "Unveiling the small protein landscape of uropathogenic *E. coli*"

**Supporting information**

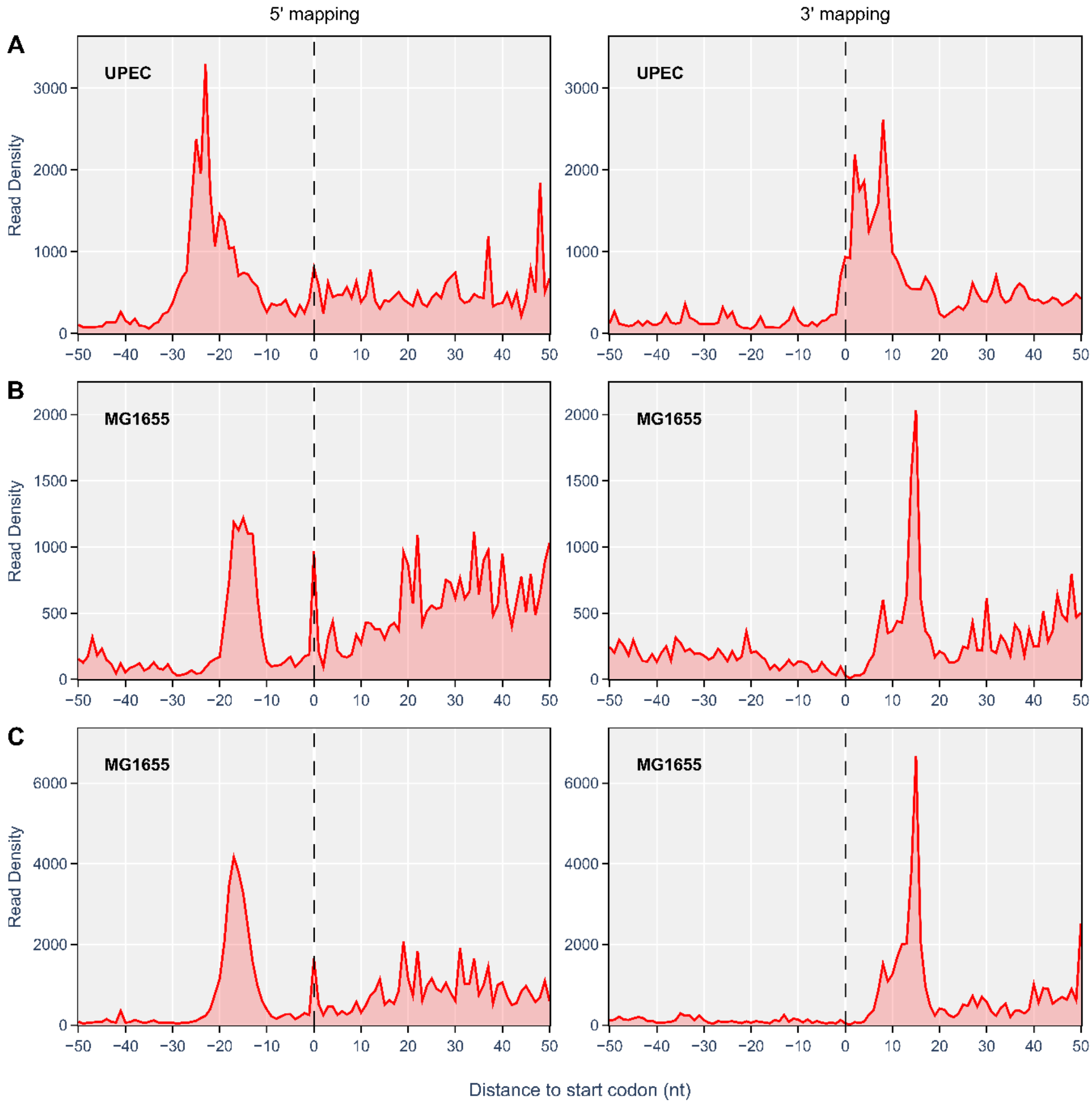


**Fig. S1. Metagene analysis of ribosome profiling datasets** *(A)* Metagene analysis showing the 3' (left) and 5' (right) ribosome footprints near the start codon of ORFs in UPEC. The same analysis was performed using published *E. coli* MG1655 ribosome profiling datasets from (B) the Storz (doi.org/10.1128/mbio.02819-18) and (C) the Buskirk laboratories (doi.org/10.1016/j.celrep.2015.03.014). The start codon locations are indicated by vertical dashed lines at 0. All plots were generated by Plotly.js (v2.9.0).

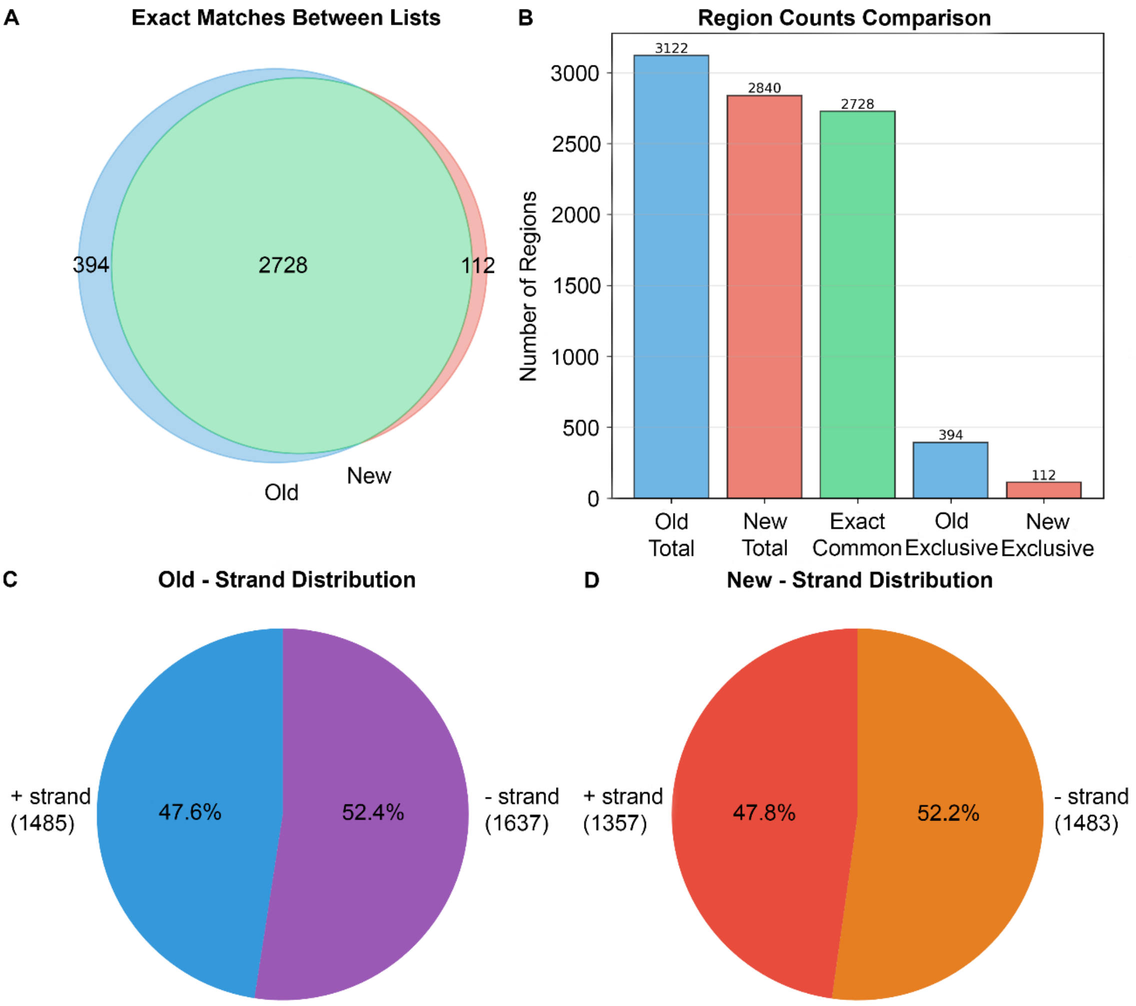


**Fig. S2. Comparison of the two rounds of HRIBO analyses (old and new) with the same set of experimental data.** Identified ORFs (A) and region counts (B) were compared. Strand distributions of two rounds of analyses are shown in C and D.

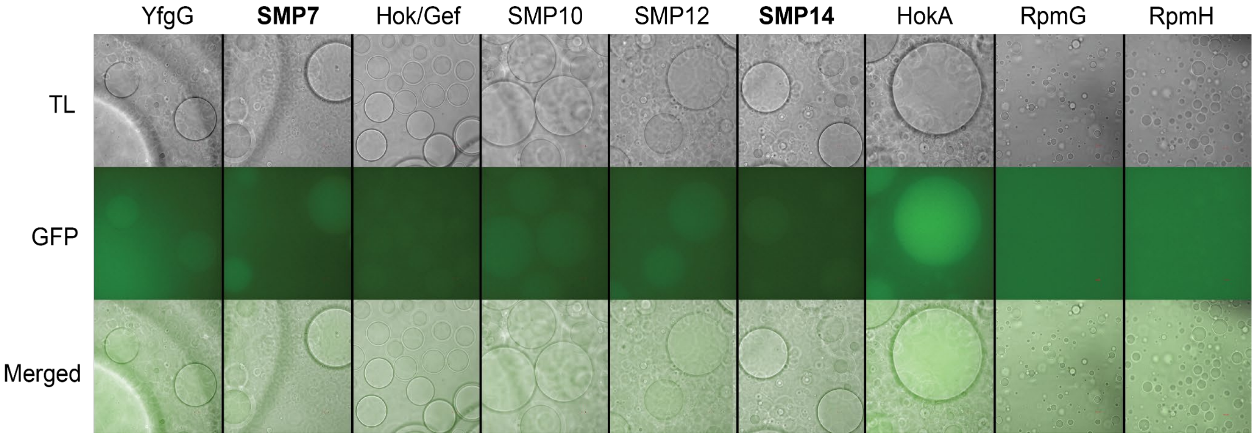


**Fig. S3. Cell-free synthesis of putative SMPs with lipid sponge droplets.** The mNeonGreen labeled SMPs were synthesized in the cell-free system containing lipid droplets *as described in Fig. 4.* Data are representative of at least two independent experiments.

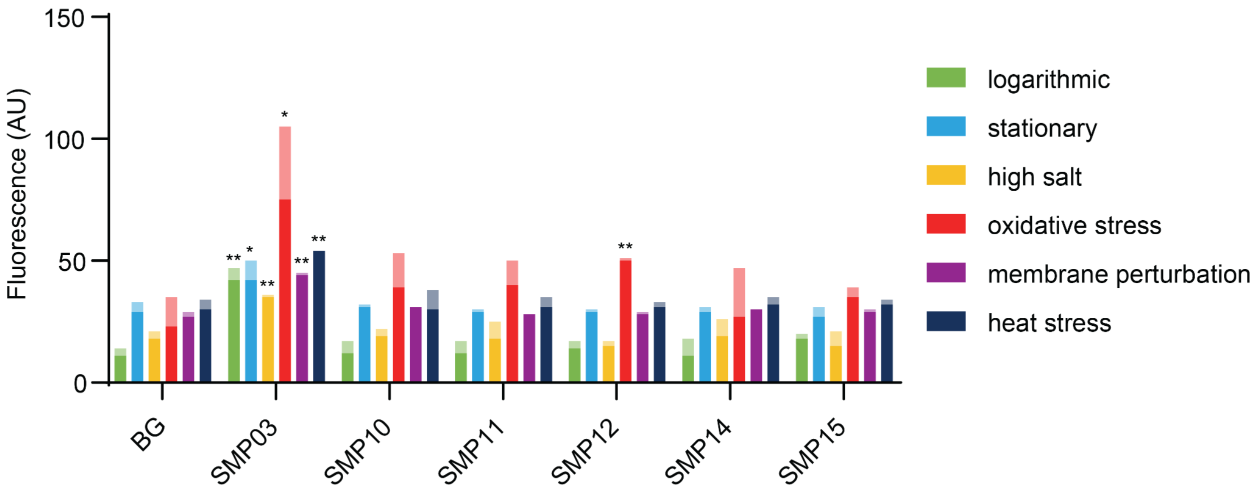


**Fig. S4. Flow cytometry–based quantification of small protein expression under the indicated stress conditions.** *Stress treatments were applied during logarithmic growth, except for heat stress, which was applied from the start of the culture. Bars with different shading denote two independent biological replicates. BG stands for background control.*

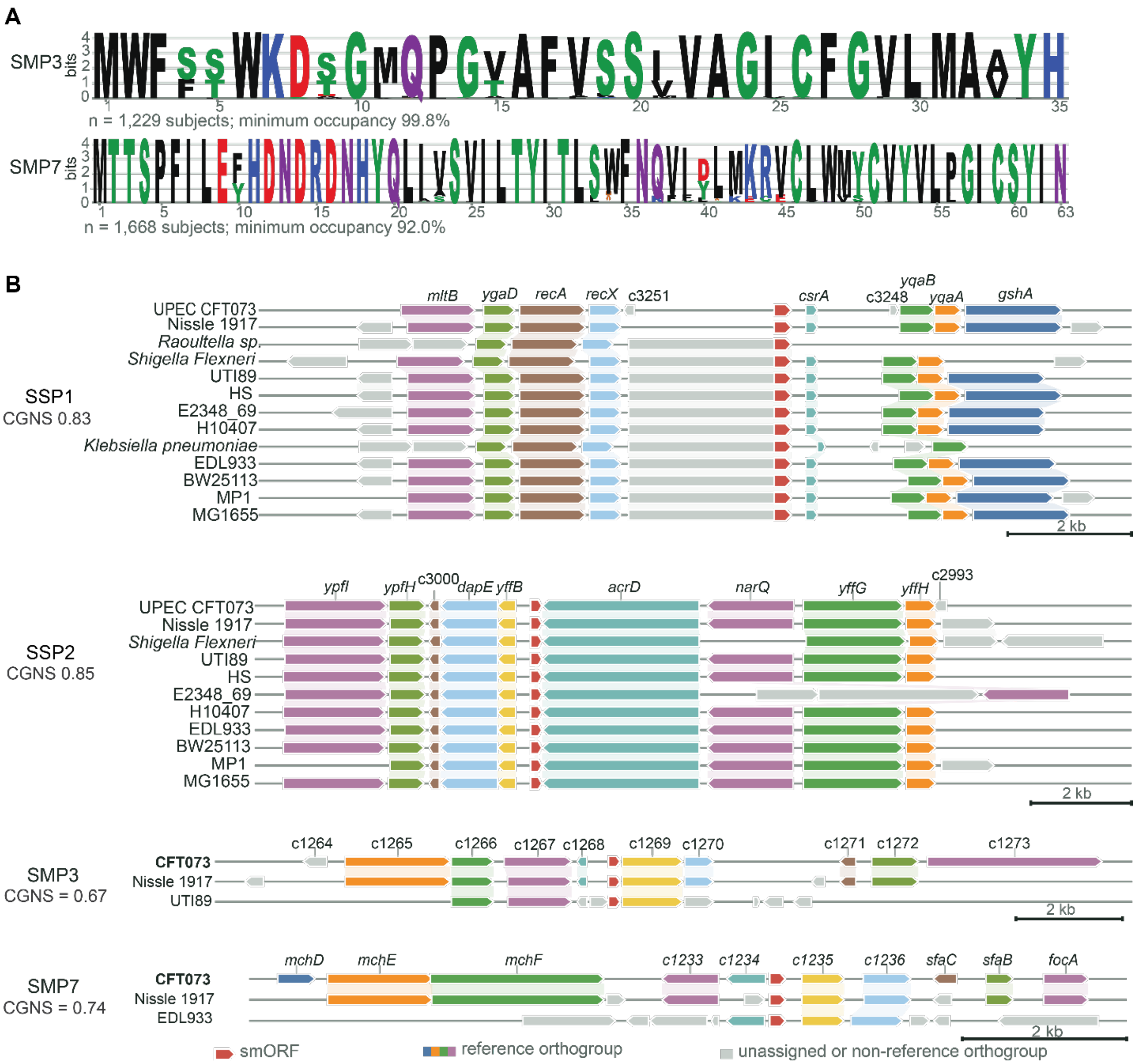


**Fig. S5. The sequence logo of putative small proteins and the genomic neighborhood of sORFs.** (**A**) The sequence logo of the translated TBLASTN hits represents the frequency of amino acids at each residue of SMP3 and SMP7. (**B**) Homologous genomic neighborhood of SSP1, SSP2, SMP3 and SMP7 is presented in the indicated genomes. The degree of conservation is indicated with the mean CGNS values.

**Table S1. List of primers used for cell-free constructs**

| Primer | 5' to 3' sequence |
|---|---|
| Common rv2 | CAAAAAACCCCTCAAGACCCGTTTAGAGGCCCCAAGGGGTTATGCTAGTCACTTGTACA ATTCGTCCATACCCAT |
| *yfgG* fw1 | GAAATTAATACGACTCACTATAGGGGAATTGTGAGCGGATAACAATTCCCCTGTAGAAA TAATTTTGTTTAACTTTAATAAGGAGATATACC ATGAGTCAGGCAACCAGTATGCGA |
| *yfgG* fw2 | GTCGAATCACCGGTACAACGTGGCGGTGGCGGTAGCATGGTTTCTAAGGGTGAAGAA |
| *yfgG* rv1 | TTCTTCACCCTTAGAAACCATGCTACCGCCACCGCCACGTTGTACCGGTGATTCGAC |
| *mgtS* fw1 | GAAATTAATACGACTCACTATAGGGGAATTGTGAGCGGATAACAATTCCCCTGTAGAAA TAATTTTGTTTAACTTTAATAAGGAGATATACC ATGCTGGGTAATATGAATGTTTTT |
| *mgtS* fw2 | GCCACAAATGGGATGACGGCGGTGGCGGTAGC ATGGTTTCTAAGGGTGAAGAA |
| *mgtS* rv1 | TTCTTCACCCTTAGAAACCATGCTACCGCCACCGCCGTCATCCCATTTGTGGC |
| *SMP3* fw1 | GAAATTAATACGACTCACTATAGGGGAATTGTGAGCGGATAACAATTCCCCTGTAGAAAT AATTTTGTTTAACTTTAATAAGGAGATATACC ATGTGGTTTTCTTCATGGAAAGAC |
| *SMP3* fw2 | CTCATGGCTGTGTACCATGGCGGTGGCGGTAGC ATGGTTTCTAAGGGTGAAGAA |
| *SMP3* rv1 | TTCTTCACCCTTAGAAACCATGCTACCGCCACCGCCATGGTACACAGCCATGAG |
| *pmrR* fw1 | GAAATTAATACGACTCACTATAGGGGAATTGTGAGCGGATAACAATTCCCCTGTAGAAAT AATTTTGTTTAACTTTAATAAGGAGATATACC ATGAAAAACCGTGTTTATGAAAGT |
| *pmrR* fw2 | TGGTTTGCCACGTACGGCGGTGGCGGTAGC ATGGTTTCTAAGGGTGAAGAA |
| *pmrR* rv1 | TTCTTCACCCTTAGAAACCATGCTACCGCCACCGCCGTACGTGGCAAACCA |
| *ynhF* fw1 | GAAATTAATACGACTCACTATAGGGGAATTGTGAGCGGATAACAATTCCCCTGTAGAAAT AATTTTGTTTAACTTTAATAAGGAGATATACC ATGAGCACCGACCTTAAATTTTCA |
| *ynhF* fw2 | CTGCCGCGCTGCACGGCGGTGGCGGTAGC ATGGTTTCTAAGGGTGAAGAA |
| *ynhF* rv1 | TTCTTCACCCTTAGAAACCATGCTACCGCCACCGCCGTGCAGCGCGGCAG |
| *SMP7* fw1 | GAAATTAATACGACTCACTATAGGGGAATTGTGAGCGGATAACAATTCCCCTGTAGAAA TAATTTTGTTTAACTTTAATAAGGAGATATACCATGACTACTTCTCCTTTCATTCTC |
| *SMP7* fw2 | CCGGGCATCTGTTCTTACATTAATGGTGGCGGTAGCATGGTTTCTAAGGGTGAAGAA |
| *SMP7* rv1 | TTCTTCACCCTTAGAAACCATGCTACCGCCACCATTAATGTAAGAACAGATGCCCGG |
| *hok/gef* fw1 | GAAATTAATACGACTCACTATAGGGGAATTGTGAGCGGATAACAATTCCCCTGTAGAAAT AATTTTGTTTAACTTTAATAAGGAGATATACCATGCTGACAAAATATGCCCTTGTG |
| *hok/gef* fw2 | AGTTCTCGCTTACGAATCGAAGAAGGGTGGCGGTAGCATGGTTTCTAAGGGTGAAGAA |
| *hok/gef* rv1 | TTCTTCACCCTTAGAAACCATGCTACCGCCACCCTTCTTCGATTCGTAAGCGAGAACT |
| *SMP10* fw1 | GAAATTAATACGACTCACTATAGGGGAATTGTGAGCGGATAACAATTCCCCTGTAGAAAT AATTTTGTTTAACTTTAATAAGGAGATATACCATGCAGGCACGGAAGCGCGTTATA |
| *SMP10* fw2 | GAAATTATGGTGTCGGCAGGTGGTGGCGGTAGCATGGTTTCTAAGGGTGAAGAA |
| *SMP10* rv1 | TTCTTCACCCTTAGAAACCATGCTACCGCCACCACCTGCCGACACCATAATTTC |
| *SMP11* fw1 | GAAATTAATACGACTCACTATAGGGGAATTGTGAGCGGATAACAATTCCCCTGTAGAAAT AATTTTGTTTAACTTTAATAAGGAGATATACCATGGGTGAGATACGTGCATCAATA |
| *SMP11* fw2 | TTCTCTTTTCGAGTCCCAGCCAGTTTGGTGGCGGTAGCATGGTTTCTAAGGGTGAAGAA |
| *SMP11* rv1 | TTCTTCACCCTTAGAAACCATGCTACCGCCACCAAACTGGCTGGGACTCGAAAAGAGAA |
| *SMP12* fw1 | GAAATTAATACGACTCACTATAGGGGAATTGTGAGCGGATAACAATTCCCCTGTAGAAAT AATTTTGTTTAACTTTAATAAGGAGATATACCATGATCGCCGCCAGTGCCACCAGT |
| *SMP12* fw2 | GCTCGACCTTACGCAGTCGCGGCGGTGGCGGTAGCATGGTTTCTAAGGGTGAAGAA |
| *SMP12* rv1 | TTCACCCTTAGAAACCATGCTACCGCCACCGCCGCGACTGCGTAAGGTCGAGC |
| *blr* fw1 | GAAATTAATACGACTCACTATAGGGGAATTGTGAGCGGATAACAATTCCCCTGTAGAAAT |

| | |
|---|---|
| | AATTTTGTTTAACTTTAATAAGGAGATATACCATGAATCGTCTTATTGAATTAACA |
| *blr* -fw2 | GTACTACAGTACAACACAAGGGCGGTGGCGGTAGCATGGTTTCTAAGGGTGAAGA |
| *blr* rv1 | TCTTCACCCTTAGAAACCATGCTACCGCCACCGCCCTTGTGTTGTACTGTAGTAC |
| *SMP14* fw1 | GAAATTAATACGACTCACTATAGGGGAATTGTGAGCGGATAACAATTCCCCTGTAGA AATAATTTTGTTTAACTTTAATAAGGAGATATACCATGCTGCTGAGTGGGGAAAGTGTG |
| *SMP14* fw2 | TTGCAAGGCCACACATGTATGGCGGTGGCGGTAGCATGGTTTCTAAGGGTGAAGA |
| *SMP14* rv1 | TCTTCACCCTTAGAAACCATGCTACCGCCACCGCCATACATGTGTGGCCTTGCAA |
| *SMP15* fw1 | GAAATTAATACGACTCACTATAGGGGAATTGTGAGCGGATAACAATTCCCCTGTAGAA ATAATTTTGTTTAACTTTAATAAGGAGATATACCATGTTTGGGATTATTAAGCTAAC |
| *SMP15* fw2 | CGAAAAAGACCGGAACGAAGGGCGGTGGCGGTAGCATGGTTTCTAAGGGTGAAGA |
| *SMP15* rv1 | TCTTCACCCTTAGAAACCATGCTACCGCCACCGCCCTTCGTTCCGGTCTTTTTCG |
| *hokA* fw1 | GAAATTAATACGACTCACTATAGGGGAATTGTGAGCGGATAACAATTCCCCTGTAGAAA TAATTTTGTTTAACTTTAATAAGGAGATATACCATGCATAGACACCATAAGGTTAGC |
| *hokA* fw2 | TTAGCCTGCAAATTGAAAGAGGGCGGTGGCGGTAGC ATGGTTTCTAAGGGTGAAGAA |
| *hokA* rv1 | TTCTTCACCCTTAGAAACCATGCTACCGCCACCGCCCTCTTTCAATTTGCAGGCTAA |
| *rpmG* fw1 | GAAATTAATACGACTCACTATAGGGGAATTGTGAGCGGATAACAATTCCCCTGTAGAAA TAATTTTGTTTAACTTTAATAAGGAGATATACCATGGCTAAAGGTATTCGTGAGAAA |
| rpmG fw2 | AAAGAAGCGAAAATCAAAGGCGGTGGCGGTAGCATGGTTTCTAAGGGTGAAGAA |
| *rpmG* rv1 | TTCTTCACCCTTAGAAACCATGCTACCGCCACCGCCCTTAGAAACGGTCAGACGAGC |
| *rpmH* fw1 | GAAATTAATACGACTCACTATAGGGGAATTGTGAGCGGATAACAATTCCCCTGTAGAAAT AATTTTGTTTAACTTTAATAAGGAGATATACCATGAAACGCACTTTTCAACCGTCT |
| *rpmH* fw2 | TCGTCTGACCGTTTCTAAGGGCGGTGGCGGTAGCATGGTTTCTAAGGGTGAAGAA |
| *rpmH* rv1 | TTCTTCACCCTTAGAAACCATGCTACCGCCACCGCCCTTAGAAACGGTCAGACGAGC |

**Table S2. List of primers for constructing *in vivo* expression plasmids**

| | |
|---|---|
| Lisa_372_Gibson_pLW 73_i_fw | GCGGTATTTCACACCGCATATGTCTAGAGTCGACGGCATGG |
| Lisa_373_Gibson_pLW 73_i_rv | CATGATCTTTGTAGTCGCCGTCGTGGTCTTTGTAGTCGCTGCCACCGCCACCCTTGT ACAATTCGTCCATAC |
| Lisa_374_Gibson_pLW 73_v_fw | CACGACGGCGACTACAAAGATCATGACATTGATTACAAAGATGATGATGATAAGT AGCTGCAGGAGAAGATTTTC |
| Lisa_375_Gibson_pLW 73_v_rv | CCATGCCGTCGACTCTAGACATATGCGGTGTGAAATACCGC |
| Lisa_376_Gibson_pLW 74_i_fw | GTATTTCACACCGCATATGTTTAAGTCGAACACAATAAG |
| Lisa_377_Gibson_pLW 74_i_rv | CCATGCCGTCGACTCTAGAGTCATCCCATTTGTGGCTG |
| Lisa_378_Gibson_pLW 74_v_fw | CAGCCACAAATGGGATGACTCTAGAGTCGACGGCATGG |

| | |
|---|---|
| Lisa_379_Gibson_pLW74_v_rv | CTTATTGTGTTCGACTTAAACATATGCGGTGTGAAATAC |
| Lisa_380_Gibson_pLW75_i_fw | GGTATTTCACACCGCATATGGAAGCTGGCCCCTGTGG |
| Lisa_381_Gibson_pLW75_i_rv | CCATGCCGTCGACTCTAGAATGGTACACAGCCATGAGAAC |
| Lisa_382_Gibson_pLW75_v_fw | GTTCTCATGGCTGTGTACCATTCTAGAGTCGACGGCATGG |
| Lisa_383_Gibson_pLW75_v_rv | CCACAGGGGCCAGCTTCCATATGCGGTGTGAAATACC |
| Lisa_384_Gibson_pLW76_i_fw | GGTATTTCACACCGCATATGCACTCGCTCAATCAACTACG |
| Lisa_385_Gibson_pLW76_i_rv | CCATGCCGTCGACTCTAGAGCCAGCACGCAAAATAAAG |
| Lisa_386_Gibson_pLW76_v_fw | CTTTATTTTGCGTGCTGGCTCTAGAGTCGACGGCATGG |
| Lisa_387_Gibson_pLW76_v_rv | CGTAGTTGATTGAGCGAGTGCATATGCGGTGTGAAATACC |
| Lisa_388_Gibson_pLW77_i_fw | GGTATTTCACACCGCATATGTATCCTTTACCCTCGCTC |
| Lisa_389_Gibson_pLW77_i_rv | CCATGCCGTCGACTCTAGACTTGTGTTGTACTGTAGTAC |
| Lisa_390_Gibson_pLW77_v_fw | GTACTACAGTACAACACAAGTCTAGAGTCGACGGCATGG |
| Lisa_391_Gibson_pLW77_v_rv | GAGCGAGGGTAAAGGATACATATGCGGTGTGAAATACC |
| Lisa_392_Gibson_pLW78_i_fw | GTATTTCACACCGCATATGGATAAATGAGACGCCAGCAC |
| Lisa_393_Gibson_pLW78_i_rv | CCATGCCGTCGACTCTAGAATTAATGTAAGAACAGATG |
| Lisa_394_Gibson_pLW78_v_fw | CATCTGTTCTTACATTAATTCTAGAGTCGACGGCATGG |
| Lisa_395_Gibson_pLW78_v_rv | GTGCTGGCGTCTCATTTATCCATATGCGGTGTGAAATAC |
| Lisa_396_Gibson_pLW79_i_fw | GTATTTCACACCGCATATGGACCCGTCGTCAAGGCGCTG |
| Lisa_397_Gibson_pLW79_i_rv | CATGCCGTCGACTCTAGATGCCGACACCATAATTTCGTC |
| Lisa_398_Gibson_pLW79_v_fw | GACGAAATTATGGTGTCGGCATCTAGAGTCGACGGCATG |
| Lisa_399_Gibson_pLW79_v_rv | CAGCGCCTTGACGACGGGTCCATATGCGGTGTGAAATAC |
| Lisa_400_Gibson_pLW80_i_fw | GTATTTCACACCGCATATGGTGAAATTCTTTTGGCGTCAG |
| Lisa_401_Gibson_pLW80_i_rv | CATGCCGTCGACTCTAGAAAACTGGCTGGGACTCGAAAAG |

| | |
|---|---|
| Lisa_402_Gibson_pLW80_v_fw | CTTTTCGAGTCCCAGCCAGTTTTCTAGAGTCGACGGCATG |
| Lisa_403_Gibson_pLW80_v_rv | CTGACGCCAAAAGAATTTCACCATATGCGGTGTGAAATAC |
| Lisa_404_Gibson_pLW81_i_fw | GTATTTCACACCGCATATGTGATTCGTTAACAAATACGC |
| Lisa_405_Gibson_pLW81_i_rv | CATGCCGTCGACTCTAGAGCGACTGCGTAAGGTCGAGCTAC |
| Lisa_406_Gibson_pLW81_v_fw | GTAGCTCGACCTTACGCAGTCGCTCTAGAGTCGACGGCATG |
| Lisa_407_Gibson_pLW81_v_rv | GCGTATTTGTTAACGAATCACATATGCGGTGTGAAATAC |
| Lisa_408_Gibson_pLW82_i_fw | GTATTTCACACCGCATATGCAAGTAATACGGCGAGCGATATG |
| Lisa_409_Gibson_pLW82_i_rv | CATGCCGTCGACTCTAGAATACATGTGTGGCCTTGCAAGC |
| Lisa_410_Gibson_pLW82_v_fw | GCTTGCAAGGCCACACATGTATTCTAGAGTCGACGGCATG |
| Lisa_411_Gibson_pLW82_v_rv | CATATCGCTCGCCGTATTACTTGCATATGCGGTGTGAAATAC |
| Lisa_412_Gibson_pLW83_i_fw | GTATTTCACACCGCATATGCATATTTCGCATTTGAATATTG |
| Lisa_413_Gibson_pLW83_i_rv | CATGCCGTCGACTCTAGACTTCGTTCCGGTCTTTTTCG |
| Lisa_414_Gibson_pLW83_v_fw | CGAAAAAGACCGGAACGAAGTCTAGAGTCGACGGCATG |
| Lisa_415_Gibson_pLW83_v_rv | CAATATTCAAATGCGAAATATGCATATGCGGTGTGAAATAC |

**Table S3. List of plasmids**

| | |
|---|---|
| pLW73 | pTrc99a_noprom_mNeonGreen-3xFlag |
| pLW74 | pTrc99a_Pnsp02-nsp02-mNeonGreen-3xFlag |
| pLW75 | pTrc99a_Pnsp03-nsp03-mNeonGreen-3xFlag |
| pLW76 | pTrc99a_Pnsp08-nsp08-mNeonGreen-3xFlag |
| pLW77 | pTrc99a_Pnsp13-nsp13-mNeonGreen-3xFlag |
| pLW78 | pTrc99a_Pnsp07-nsp07-mNeonGreen-3xFlag |
| pLW79 | pTrc99a_Pnsp10-nsp10-mNeonGreen-3xFlag |
| pLW80 | pTrc99a_Pnsp11-nsp11-mNeonGreen-3xFlag |
| pLW81 | pTrc99a_Pnsp12-nsp12-mNeonGreen-3xFlag |
| pLW82 | pTrc99a_Pnsp14-nsp14-mNeonGreen-3xFlag |
| pLW83 | pTrc99a_Pnsp15-nsp15-mNeonGreen-3xFlag |

**Table S4. List of formulae used for conserved**

| Component | Template | Formula | Definition |
|---|---|---|---|
| **Presence** | consensus or CFT073 | $P_s(T) = \frac{m_s(T)}{n_T}$ | Exact positional matches to template T. |
| **Distance** | consensus or CFT073 | $D_s(T) = \frac{1}{n_T}\sum_{i \in T}\left[1 - \min\left(1, \frac{d_{i,s}(T)}{2W+1}\right)\right]$ | Nearest-position recovery of template orthogroup i; W = 5. |
| **Order** | consensus or CFT073 | $O_s(T) = \frac{c_s(T)}{r_s(T)}$ | Concordant relative order among mapped template-orthogroup pairs. |
| **Species CGNS** | consensus or CFT073 | $\mathrm{CGNS}_s(T) = 0.5P_s(T) + 0.3O_s(T) + 0.2D_s(T)$ | Species-level score against the selected template. |
| **Consensus mean** | consensus | $\overline{\mathrm{CGNS}}_{\mathrm{cons}} = \frac{1}{n}\sum_{s}\mathrm{CGNS}_s(T_{\mathrm{cons}})$ | Main aggregate; the E. coli mean restricts s to the predefined E. coli subset. |
| **CFT073-reference mean** | CFT073 | $\overline{\mathrm{CGNS}}_{\mathrm{CFT073}} = \frac{1}{n_{\mathrm{ref}}}\sum_{s \neq \mathrm{CFT073}}\mathrm{CGNS}_s(T_{\mathrm{CFT}})$ | Reference-template check, excluding the CFT073 self-score. |
